# Restricted intake of sulfur-containing amino acids reversed the hepatic injury induced by excess *Desulfovibrio* through gut-liver axis

**DOI:** 10.1101/2024.02.01.578326

**Authors:** Lingxi Zhou, Gexue Lu, Yawen Nie, Yilin Ren, Jin-Song Shi, Yuzheng Xue, Zheng-Hong Xu, Yan Geng

## Abstract

Gut-liver axis has been a study focus for liver diseases. Diet is a key player in influencing the gut microbiota. However, the effect of different dietary patterns on gut microbiota and liver functions remains unclear. Here, we used mouse standard chow and purified diet to mimic two common human dietary patterns: healthy grain and planted-based diet and Western style diet, respectively and explored their impacts on the gut microbiota and liver. Gut microbiota experienced a great shift with notable increase in *Desulfovibrio*, gut bile acid (BA) concentration elevated significantly, and liver inflammation was observed in mice fed with the purified diet. Liver inflammation due to translocation of toxic lipopolysaccharides (LPS) and hydrophobic BAs from the damaged gut barrier was also observed in mice fed with the chow diet after receiving *Desulfovibrio desulfuricans* ATCC 29577 (DSV). Restricted intake of sulfur-containing amino acids reversed the liver injury due to excess *Desulfovibrio* through lowering the gut BA concentration and enhancing the hepatic antioxidant and detoxifying ability. *Ex vivo* fermentation of human fecal microbiota with primary BAs also demonstrated that DSV enhanced production of secondary BAs. Germ-free mice had higher concentration of both conjugated and unconjugated primary BAs in their gut after receiving DSV.

## Introduction

The concept of liver-gut axis was firstly proposed by Marshall in 1998 and has been studied for over 20 years. Understanding the crosstalk between the liver and gut helps elucidate many causes behind the liver diseases on the liver disease spectrum from liver inflammation at the beginning to gradually advancing diseases, including non-alcoholic fatty liver disease (NAFLD), alcoholic liver disease (ALD), liver fibrosis, liver cirrhosis and liver cancer in the end^1, 2^. The liver is affected by the gut when nutrients, microbial metabolites and microbial particles are transferred from the gut to liver through the portal vein; on the other hand, gut microbial growth and metabolism are closely related to bile acids secreted by the liver^1–5^. When gut dysbiosis occurs, impaired gut barrier allows translocation of potentially pathogenic microbes, harmful microbial metabolites and toxins to the liver, which makes the liver overwhelmed by pro-inflammatory molecules and causes liver diseases ultimately if no interventions are involved^6–10^.

Dietary factors always play an essential role in the gut-liver axis since food is the most active interactor with gut microbes, which pose a direct impact on the microbial structure and microbial metabolism. Many researches have focused on how dietary patterns affect host health through gut-host interaction. Unhealthy diets such as high-fat diet has always been the study focus for metabolic diseases including obesity, diabetes, metabolic liver diseases, cardiovascular diseases and so on while healthy diets such as Mediterranean diet and methionine-restricted (MR) diet are associated with distinct gut microbiota, healthy gut barrier, improved host metabolism and longevity^1, 11–13^. MR diet, as a type of sulfur-containing-amino-acid-limited (SAAL) diet, has been studied as a dietary strategy for longevity which could extend health span and lifespan in rodents by improving the glucose metabolism, enhancing antioxidant ability to liver cell injury, reducing energy expenditure, reversing the transcriptional alterations in inflammation and DNA-damage response genes, reducing hepatic DNA hypermethylation and improving the lipid and BA profile in aging adult mice or progeria mouse models^13–16^.

The chow diet and purified diet are widely used in rodent studies, which can mimic two common dietary patterns of humans: healthy grain and planted-based diet with minimal processing and Western style diet made of more refined and over-processed ingredients, respectively. The biggest difference between the chow and purified diet is the degree of refinement of ingredients and the bioavailability of nutritional contents. Chow diet is mainly made of natural ingredients such as whole grains, high protein meals, together with some vitamin and mineral premixes. The nutrients are often in bound forms in chow diet and can be variable from batch to batch. Purified diet is often made of more refined ingredients such as casein, corn starch, maltodextrin, vegetable oil and mixed with chemically pure inorganic salts and vitamins, which are more chemically and nutritionally defined^17^. Since purified diet is more refined and contains limited types of ingredients, the nutrients are more accessible in unbound forms and at the same time lack some bioactive compounds existing in unprocessed food ingredients, which may pose unpredictable health effects on the host. Recent studies demonstrate that compared with the chow diet, the purified diet is more disease-inducing, including altered gut microbial structure, increased gut barrier damage, increased hepatic steatosis and inflammation, impaired glucose tolerance, and exacerbation of DSS-induced colitis and inflammatory bowel disease (IBD) even supplemented with commonly recognized beneficial oligosaccharides or fibers^18–24^. However, how the purified diet modulates the gut microbiota and thus affects liver pathophysiology is poorly understood.

In this study, we aim to investigate the impacts and underlying mechanism of dietary patterns on gut microbiota and liver health. We confirmed the disease-inducing nature of the purified diet, in which gut microbiota and microbial metabolic profile experience a great shift and liver inflammation is observed. We then propose and validate that *Desulfovibrio*, a key microbial biomarker in mice fed with the purified diet, as an important player in the alteration of gut microbial BA metabolism and liver inflammation. We further extend the health effects of SAAL diet, where it can reverse the hepatic injury and elevation of gut BA concentration due to extra *Desulfovibrio* in mice. Overall, this study provides new insights of how dietary patterns affect liver health by proposing *Desulfovibrio* as a key microbial player. Restricted intake of sulfur-containing amino acids may become an effective dietary intervention to reverse the liver injury associated with excess *Desulfovibrio* in the gut.

## Materials and methods

### Animals and treatments

5-week-old male BALB/c mice were purchased from Shanghai Slac Laboratory Animal Ltd. (Shanghai, China). Mice were housed in a specific-pathogen free (SPF) environment (20-26 ℃, 40-70% humidity, 12 light/dark cycle) with ad libitum access to distilled water and sterilized food.

#### 1st animal experiment

After one-week acclimation, mice were randomly divided into 2 groups according to their body weight (n = 10 mice/group). Mice in the CTRL group were fed with the chow diet (M01-F, Shanghai Slac Laboratory Animal Ltd., Shanghai, China) and mice in the PURE group were fed with the purified diet (TP0016S, Trophic Animal Feed High-Tech Co. Ltd., China). This purified diet is an amino-acid defined NRC95 standard diet. NRC95 refers to the fourth revised edition of *Nutritional Requirements for Laboratory Animals* produced by National Research Council (US) Subcommittee on Laboratory Animal Nutrition in 1995, which raised the content of methionine, vitamin K and vitamin B12^17^. Amino acid-defined NRC95 purified diet refers to purified diet made according to the NRC95 standard but individual amino acids instead of casein are added. It is usually used as the control diet for animal studies which involve in change of contents of amino acids. The experiment duration was 28 days.

#### 2nd animal experiment

After one-week acclimation, mice were randomly divided into 2 groups according to their body weight (n = 5 mice/group). Mice in the CTRL group were fed with the chow diet (M01-F, Shanghai Slac Laboratory Animal Ltd., Shanghai, China) and mice in the CTRL_D group were fed with the same chow diet and administrated with 5×10^7^ CFU of DSV by oral gavage every day. DSV were purchased from China General Microbiological Culture Collection Center (CGMCC). The experiment duration was 28 days.

#### 3rd animal experiment

After one-week acclimation, mice were randomly divided into 4 groups according to their body weight (n = 6 mice/group). Mice in the PURE group were fed with the purified diet (TP0016S, Trophic Animal Feed High-tech Co. Ltd., Nantong, China). Mice in the SAAL group were fed with the SAAL diet (TP0030, containing 0.15% methionine and 0% cystine, Trophic Animal Feed High-tech Co. Ltd., Nantong, China). Mice in the PURE_D group were fed with the same diet as the PURE group and administrated with 5×10^7^ CFU of DSV by oral gavage every day. Mice in the SAAL_D group were fed with the same diet as the SAAL group and administrated with 5×10^7^ CFU of DSV by oral gavage every day. The experiment duration was 28 days.

#### 4th animal experiment

Germ-free 5-week-old BALB/c mice were bred and maintained in special plastic isolators (GemPharmatech, Nanjing, China). All mice were housed under a strict 12-hour light cycle (lights on at 08:00). Animals were supplied with a 50-kGy irradiated sterile pelleted normal chow diet (1010063, Jiangsu Xietong Pharmaceutical Bio-engineering Co., Ltd., Nanjing, China) and autoclaved tap water ab libitum. Bedding was replaced in all experiments every 7 days. All germ-free mice were routinely screened for bacteria, viral, and fungus contamination. After one-week acclimation, mice were randomly divided into 2 groups according to their body weight (n = 5 mice/group). Mice in the GFC group were fed with the chow diet and mice in the GFDSV group were fed with the chow diet and administrated with 2×10^8^ CFU of DSV by oral gavage every day for 14 days.

Study protocols comformed to the Animal research: Reporting of In Vivo Experiments (ARRIVE) guidelines. All procedures were approved by the Institutional Animal Care and Use Committee of Jiangnan University, Wuxi, China (Approval No. 20210530b0800731[137]).

### Statistical analysis

GraphPad Prism (v8.4.2, San Diego, CA) was used for data analysis. Student t-test was used for data with normal distribution and Mann-Whitney test was used for data not conforming to normal distribution for comparisons between two groups. One-way ANOVA with Holm-Sidak or FDR (Two-stage step-up method of Benjamini, Krieger and Yekutieli) post hoc test was performed for data with normal distribution and Kruskal-Wallis test with Dunn or FDR post hoc test (Two-stage step-up method of Benjamini, Krieger and Yekutieli) was performed for data not conforming to normal distribution for comparisons between three or more groups. For *ex vivo* experiment of bile acid metabolism of human fecal bacteria, data were analyzed using paired t-test or Wilcoxon matched-pairs signed rank test. Data was represented by mean ± SEM. Biological significance was represented by \**p* < 0.05 and \*\**p* < 0.01.

Supplementary methods provide additional details on sample collection, physiological parameters, histological examination, 16s rRNA gene sequencing, metagenomic sequencing, metabolomic study, transcriptomic study, gene expressions determined by qRT-PCR, evaluation of heaptic oxidative stress, *ex vivo* fermentation of human fecal bacteria with bile acids, quantification of BAs by UPLC-MS-MS and detection of amino acids in the chow diet using HPLC.

## Results

### Purified diet led to significant changes of the structure and metabolism of gut microbiota

To investigate the impact of two dietary patterns on gut microbial structure, we first fed mice in the CTRL group with chow diet and mice in the PURE group with purified diet for 28 days (Fig. 1A; nutrition facts can be referred to Table S1).

**Fig. 1.**
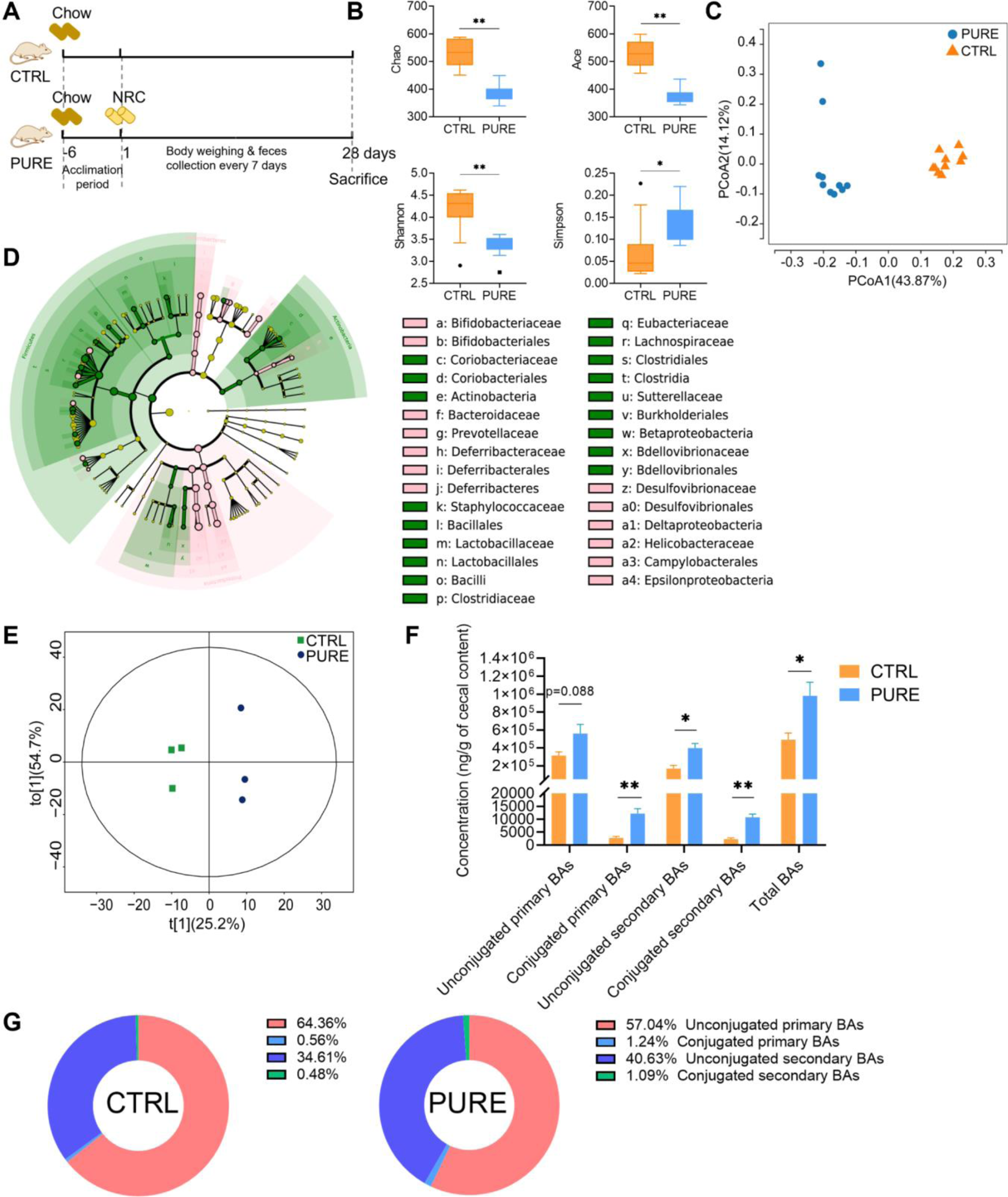
Alteration of structure and metabolism of gut microbiota in mice fed with the chow diet and purified diet. (A) Experimental design. (B) Indices of alpha diversity including Chao, Ace, Shannon and Simpson. (C) Principle Co-ordinates Analysis (PCoA, unweighted Unifrac) of gut microbiota. (D) Cladogram of LEfSe of gut microbiota. OTUs with LDA score >2 were screened and represented the microbial biomarkers in that group. (E) Orthogonal Partial Least Squares-Discriminant Analysis (OPLS-DA) of cecal metabolites. (F) Concentrations of unconjugated primary BAs, conjugated primary BAs, unconjugated secondary BAs, conjugated secondary BAs and total BAs in the mouse ceca. (G) Percentage composition of BAs. Mean values of each BA type were used to draw the pie charts. n =10 mice per group in (A-D), n = 3 mice per group in (E-G). Data are represented as the mean ± SEM, two-tailed Student t-test or Mann-Whitney test, \**p* < 0.05, \*\**p* < 0.01.

Big difference in gut microbial structure between the two groups was observed in 16s rRNA gene sequencing analysis. Indices of alpha diversity reflected higher richness and diversity of microorganisms in the CTRL group (Fig. 1B). Principle Co-ordinates Analysis (PCoA) demonstrated distinct microbial structure between the two groups (Fig. 1C). A large number of microbial biomarkers were identified in two groups by linear discriminant analysis of effect sizes (LEfSe), in which Proteobacteria, Desulfovibrionales, Desulfovibrionaceae, *Desulfovibiro* and Deltaproteobacteria were top 5 biomarkers in the PURE group (Fig. 1D, Fig. S1).

We then performed a targeted metabolomic study of the mouse cecal content to see how the microbial metabolism changed due to different diets. In total, 650 metabolites were analyzed and 348 metabolites were detected. The microbial metabolic profiles of the CTRL and PURE group were very different (Fig. 1E, Fig. S2). Metabolites with significant difference were mainly consist of three groups: BAs, amino acids and their derivatives and carbohydrates and their derivatives. Alteration of BA profile was evident with the highest number of changed compounds. The concentrations of conjugated primary BAs, unconjugated secondary BAs and conjugated secondary BAs significantly elevated in the PURE group (Fig. 1F). Meanwhile, the percentage of unconjugated primary BA was lower while percentage of unconjugated secondary BA and conjugated secondary BA was higher in the PURE group compared with those in the CTRL group, indicating active transformation of unconjugated primary BAs to secondary BAs by gut microbes (Fig. 1G). The concentration of secondary BAs including 3-dehydrocholic acid (3-DHCA) (a.k.a. 3-oxocholic acid), 7-dehydrocholic acid (7-DHCA), 7-ketolithocholic acid (7-KLCA), deoxycholic acid (DCA), hyodeoxycholic acid (HDCA), isodeoxycholic acid (IsoDCA), lithocholic acid (LCA), omega-muricholic acid (ω-MCA), ursocholic acid (UCA), taurodeoxycholic acid (TDCA) and tauro-ω-muricholic acid (ω-TMCA) elevated remarkably in the PURE group (Fig. S3B). Moreover, the increased concentration of conjugated primary BAs in the mouse cecum indirectly reflected the enhanced host BA production in the PURE group (Fig. S3A). These results indicated that the purified diet modulated the structure of gut microbiota and BA metabolism.

### Purified diet led to compromised hepatic and gut barrier function

Purified diet led to increased abundance of *Desulfovibrio,* which is a type of gram-negative bacteria in the gut and may become an opportunistic pathogen. Previous studies have shown that increased abundance of *Desulfovibrio* in both mouse and human fecal samples is associated with different stages of hepatic carcinoma including steatosis, steatohepatitis, liver fibrosis, and liver cirrhosis^25^. Also altered microbial BA metabolism have been associated with intestinal and liver diseases^1, 2, 5, 26^. Therefore, liver and colonic functions were assessed to see if they were affected. Basic physioplogical parameters including daily energy intake, body weight and liver/body weight ratio were first compared. Though the daily energy intake of mice in the PURE group was lower than that of mice in the CTRL group, the body weight gain was higher in the PURE group, suggesting that purified diet was easier for mice to gain weight than the chow diet (Fig. S4A and C). Liver/body weight ratio decreased significantly in the PURE group, indicating smaller liver in mice fed with the purified diet (Fig. S4B). Hematoxylin and eosin staining (H&E) of liver sections demonstrated inflammatory infiltration in the PURE group (Fig. 2A). The clear signals of the F4/80 and Ly6G immunohistochemical (IHC) staining of mouse livers in the PURE group indicated accumulation of macrophages and neutrophils, respectively (Fig. 2B-C). Transcriptomic study showed very different gene expression patterns between mice in the two groups (Fig. S5A). 570 up-regulated genes and 399 down-regulated genes were found in the PURE group when comapring with the CTRL group (Fig. S5B). Gene expression levels with absolute value of log_2_ of fold change larger than 0.58, Q value (adjusted p value) smaller than 0.05 were selected to conduct KEGG pathway enrichment analysis, which were annotated to 30 pathways (Fig. 2D). Among these pathways, glutathione metabolism and bile secretion were further analyzed. Glutathione metabolism is highly involved in cytoprotection from oxidative stress generated both internally by the host metabolism and externally by xenobiotics such as heavy metals, drugs, toxins and bacterial LPS^27, 28^. GSEA demonstrated that glutathione metabolism was down-regulated in the PURE group (Fig. 2E). Gene expression of glutathione S-transferases (GSTs) including family alpha (A), mu (M), pi (P) and theta (T) were significantly down-regulated in the PURE group, which indicated reduced hepatic detoxifying and cytoprotective ability (Fig. 2F). GSEA of bile secretion metabolism was also down-regulated in the PURE group, in which the gene expression of Cytochrome P450 family 7 subfamily A member 1 (*Cyp7a1*) and Cytochrome P450 family 8 subfamily B member 1 (*Cyp8b1*) decreased and Nuclear receptor subfamily 1 group H member 4 (*Nr1h4*) increased (Fig. 2G-H). *Cyp7a1* is the rate-limiting enzyme in the classic pathway of BA synthesis, its down-regulation represents less BA production^26^. *Nr1h4* encodes Farnesoid X-Activated Receptor (FXR), whose expression inhibits *Cyp7a1* transcription^26^. Inhibition of *Cyp7a1* in the mouse liver of the PURE group is very possibly due to the negative feedback to excessive primary BAs and secondary BAs detected in the cecum which may directly activate the hepatic FXR via circulating back to the liver through the portal vein or indirectly activate the intestinal FXR and induce production of fibroblast growth factor 15 (FGF15), which could bind to the fibroblast growth factor receptor 4 (FGFR4) in the liver to inhibit BA synthesis^1^.

**Fig. 2.**
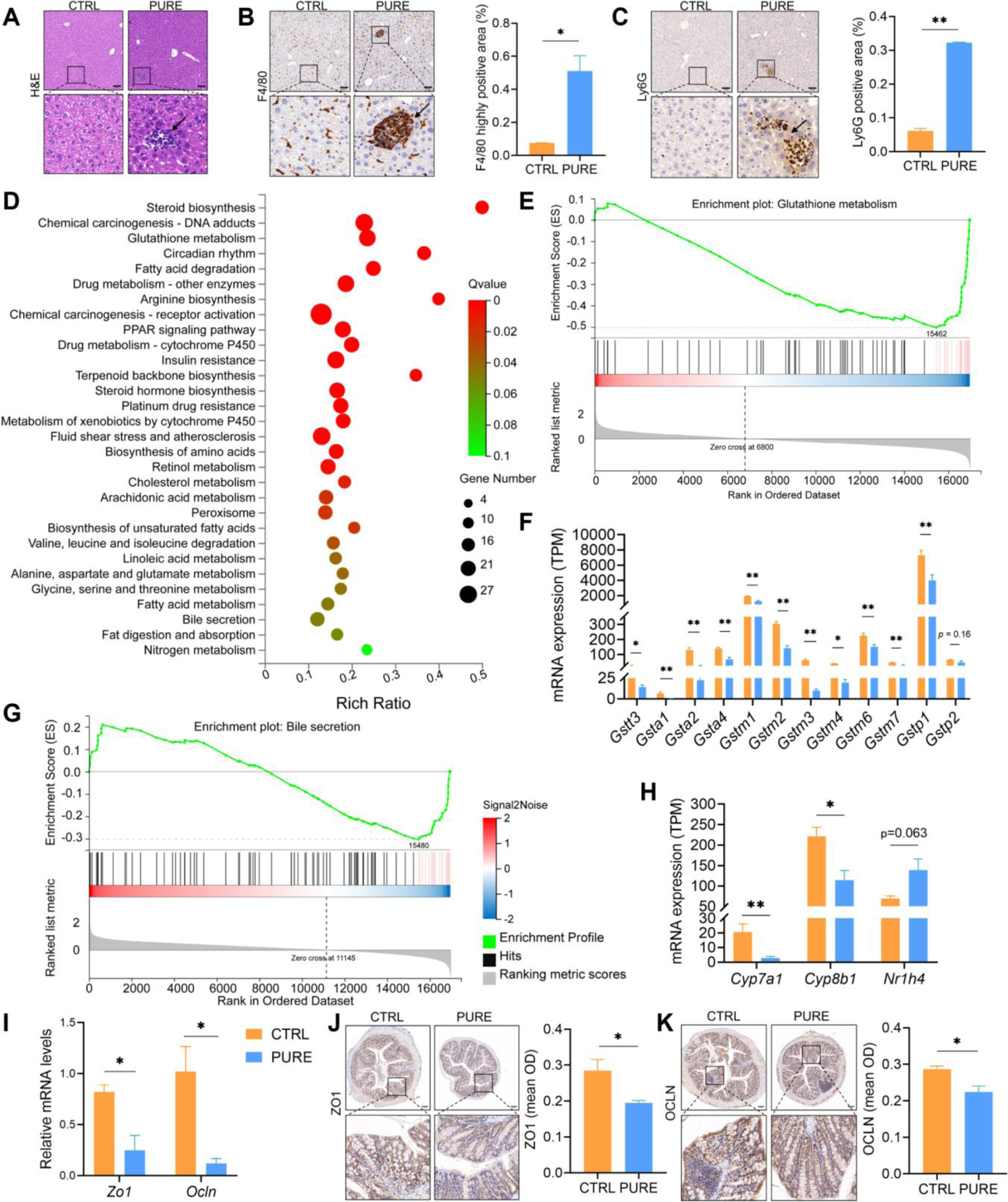
Hepatic and colonic changes of mice fed with the chow diet and purified diet. (A) H&E staining of liver sections. (B) IHC staining of F4/80 and quantification of highly positive areas of liver sections. (C) IHC staining of Ly6G and quantification of positive areas of liver sections. Black arrows point to the accumulation of immune cells. Scale bar: 100μm. (D) KEGG enrichment analysis. Gene expression levels with absolute value of log2|FC| > 0.58, Qvalue (FDR corrected p value) < 0.05 in student t-test were subject to hypergeometric test, pathway Qvalue (FDR corrected p value) < 0.1 were shown. (E) GSEA plot of glutathione metabolism when the PURE group compared with the CTRL group. The leading edge genes (red lines) are on the right, which represent up-regulation of this pathway in the CTRL group. (F) Expression levels of genes encoded Glutathione S-transferase (GST) family alpha (Gsta), mu (Gstm), pi (Gstp) and theta (Gstt). (G) GSEA plot of bile secretion pathway when the PURE group compared with the CTRL group. The leading edge genes (red lines) are on the right, which represent up-regulation of this pathway in the CTRL group. (H) Gene expression levels of hepatic *Cyp7a1*, *Cyp8b1* and *Nr1h4* in transcriptomic analysis. TPM: transcripts per million. (I) Relative mRNA expression of *Zo1* and *Ocln* in the colon using determined by qRT-PCR. (J) IHC staining of ZO1 and quantification of mean optical density (OD) of the signal. (K) IHC staining of OCLN and quantification of mean OD of the signal. Scale bar: 100μm. n = 3-5 mice per group. Data are represented as mean ± SEM, two-tailed Student t-test or Mann-Whitney test, \**p* < 0.05, \*\**p* < 0.01.

Gene expressions of Zona occludens 1 (Zo1) and Occludin (Ocln) were down-regulated in the mouse colon of the PURE group and the result of IHC staining corresponded to their gene expression levels, indicating compromised gut barrier functions (Fig. 2I-K). Excessive BAs especially the unconjugated secondary BAs (Fig. 1F) and the LPS produced by gram-negative Proteobacteria including potentially pathogenic *Desulfovibrio* and *Helicobacter* (Fig. S1) could damage the gut barrier, which allowed translocation of the toxic compounds including BAs and LPS themselves to the liver in the PURE group. BAs are naturally cytotoxic and hydrophobic BAs are usually more toxic than hydrophilic BAs^1, 26^. Unconjugated BAs are usually more hydrophobic than their corresponding conjugated BAs and secondary BAs are often more hydrophobic than their corresponding primary BAs^29^. LPS produced by gram-negative Proteobacteria are pro-inflammatory^30^. Together, these data indicate that the overwhelmed toxins together with the compromised antioxidant and detoxifying ability led to elevated hepatic oxidative stress and inflammation in mice fed with the purified diet.

### DSV administration led to hepatic injury in mice fed with the chow diet

The previous experiment suggested that purified diet could alter the gut microbiota and impact liver functions. *Desulfovibrio* was the genus that had the highest LDA score in LEfSe of gut microbiota of mice in the PURE group (Fig. S1). Previous researches have reported that the relative abundance of *Desulfovibrio* and Desulfovibrionales increase during different stages of liver diseases including hepatic steatosis, non-alcoholic steatohepatitis (NASH), hepatic fibrosis, hepatic cirrhosis and hepatic carcinoma^25, 31^. *D. desulfuricans* is a commonly found *Desulfovibrio* species^32^. To assess if *Desulfovibrio* played a key role in the alteration of microbial BA profile and liver dysfunction, a type strain of *Desulfovibrio desulfuricans* (ATCC 29577) was administrated to mice by oral gavage for 28 days (CTRL_D) and same volume of PBS was administrated to mice in the control group (CTRL) (Fig. 3A). Mice in both groups were fed with the chow diet. DSV administration decreased daily energy intake, body weight and liver/body weight ratio (Fig. S7). Accumulation of immune cells and necrosis of liver cells were observed in H&E staining in the CTRL_D group (Fig. 3B). F4/80 and Ly6G IHC staining demonstrated accumulation of macrophages and neutrophils, respectively (Fig. 3C-D). Relative gene expression levels of antioxidant and detoxifying genes including Superoxide Dismutase 1 (*Sod1*), *Sod2*, Glutathione S-transferase alpha 2 (*Gsta2*), Glutathione S-transferase Mu1 (*Gstm1*), *Gstm2* and Glutathione S-transferase pi 1 (*Gstp1*) were determined. Although the relative expression levels of *Sod1* and *Sod2* had no difference between the two groups (Fig. 3E), the SOD activity assay showed a significant reduction in the CTRL_D group, which indicated compromised antioxidant activity (Fig. 3F). Interestingly, *Gstm1*, *Gstm2* and *Gstp1* had their relative expressions enhanced in the CTRL_D group, which could be a positive response to the enhanced oxidative stress (Fig. 3E); however, the total GST activity in the CTRL_D group did not get increased (Fig. 3G). The relative expression of Acyloxyacyl hydrolase (*Aoah*) was also determined in the mouse livers to see if there was any change in the hepatic ability to detoxify LPS. AOAH is a host lipase mainly produced by macrophages, neutrophils and NK cells, which can inactivate LPS produced by gram-negative bacteria through deacylating the secondary fatty acyl chains of the lipid A in LPS and prevent further LPS - Toll-like receptor 4 (TLR4) inflammatory responses^33, 34^. The relative expression of *Aoah* in the CTRL_D group was reduced compared with that in the CTRL group (p = 0.0959), which suggests compromised LPS-detoxifying ability (Fig. 3I). Together, these results deomonstrate that *Desulfovibrio* could induce hepatic injury in mice.

**Fig. 3.**
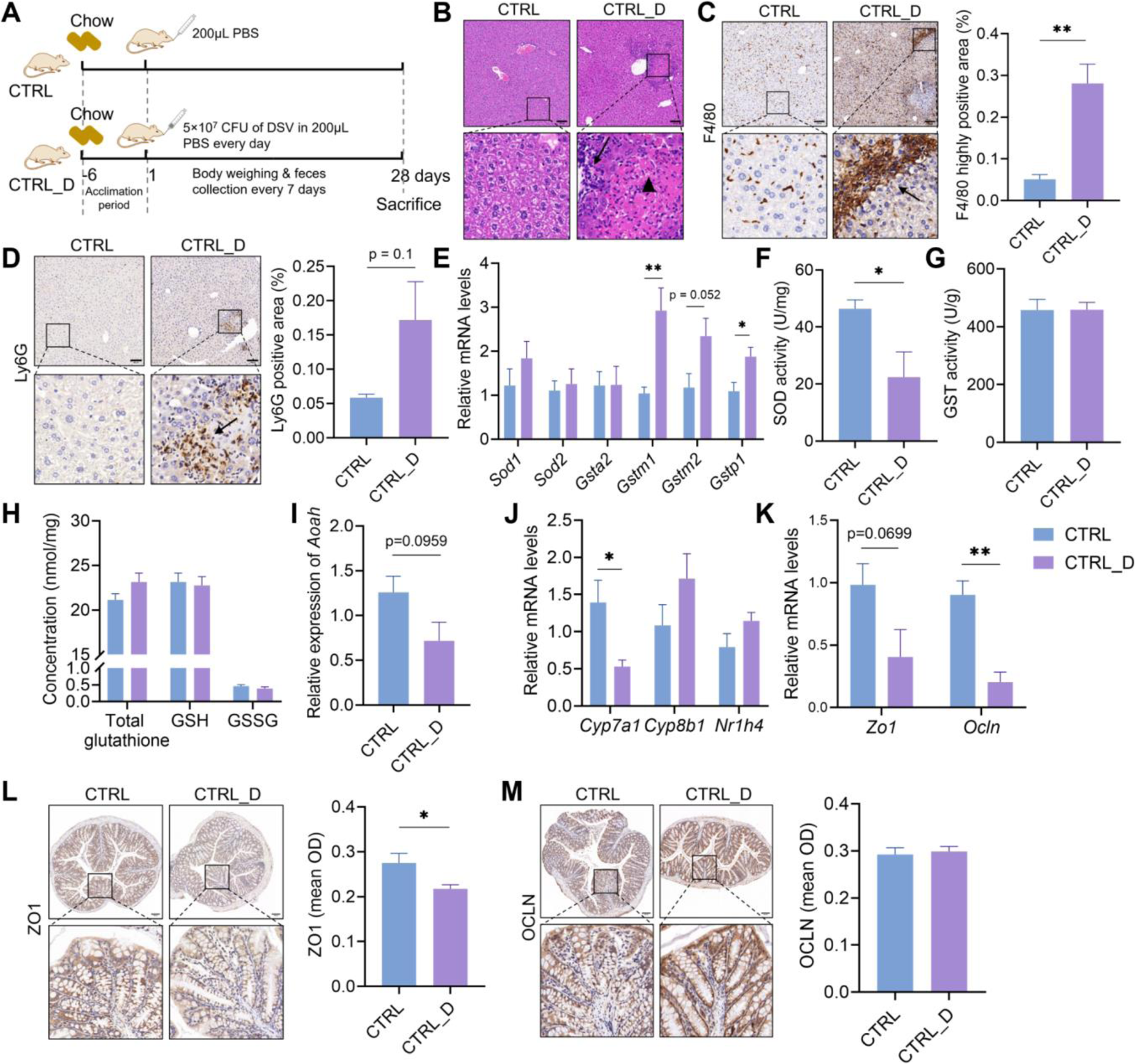
Liver and colonic changes in mice fed with the chow diet and administrated with DSV. (A) Experimental design. (B) H&E staining of liver sections. (C) IHC staining of F4/80 and quantification of highly positive areas of liver sections. (D) IHC staining of Ly6G and quantification of positive areas of liver sections. Scale bar: 100μm. Black arrows point to the accumulation of immune cells in (B-D). Black triangle indicated hepatocellular necrosis in (B). (E) Relative mRNA levels of *Sod1*, *Sod*2, *Gsta2*, *Gstm1*, *Gstm2* and *Gstp1*. *Gsta2*, *Gstm1*, *Gstm2* and *Gstp1* were selected as the representatives of GST family based on their high expression levels and big difference between the CTRL and PURE group in transcriptomic analysis in the previous experiment. (F) Hepatic SOD activity. (G) Hepatic GST activity. (H) Concentrations of hepatic total glutathione, reduced glutathione (GSH) and oxidized glutathione (GSSG). (I) Relative mRNA levels of heaptic *Aoah*. (J) Relative mRNA levels of hepatic *Cyp7a1*, *Cyp8b1* and *Nr1h4*. (K) Relative mRNA levels of *Zo1* and *Ocln* in the mouse colon. (L) IHC staining of ZO1 and quantification of mean OD of the signal. (M) IHC staining of OCLN and quantification of mean OD of the signal. Scale bar: 100μm. N = 3-5 mice per group. Data are represented as mean ± SEM, two-tailed Student t-test or Mann-Whitney test, \**p* < 0.05, \*\**p* < 0.01.

### DSV administration shifted the gut BA profile and damaged the gut barrier in mice fed with the chow diet

Metagenomic analysis of mouse cecal content were performed to analyze the difference in the gut microbial structure, which is more precise than 16s rRNA gene sequencing since it increases the sequencing depth. Chao and ace index in alpha diversity analysis showed higher richness of gut microbiota in the CTRL group than that in the CTRL_D group while the Shannon and Simpson index had no difference between the two groups (Fig. 4A). Dimensional reduction analysis could not separate the two groups (Fig. 4B). LDA score of LEfSe showed that Staphylococcaceae, *Staphylococcus*, *Ruminucoccus gnavus* and *Mediterraneibacter* were the biomarkers of the CTRL_D group while *Bacillus subtilis*, *Lactococcus* and Streptococcaceae were the biomarkers of the CTRL group (Fig. 4C). DSV administration did not increase the abundance of *Desulfovibrio* and did not alter the structure of gut microbiota too much, which is probably due to DSV’s poor colonizing ability because of low tolerance of gastric and enteric juice before reaching the colon or low competitiveness compared with the already resident commensals.

**Fig. 4.**
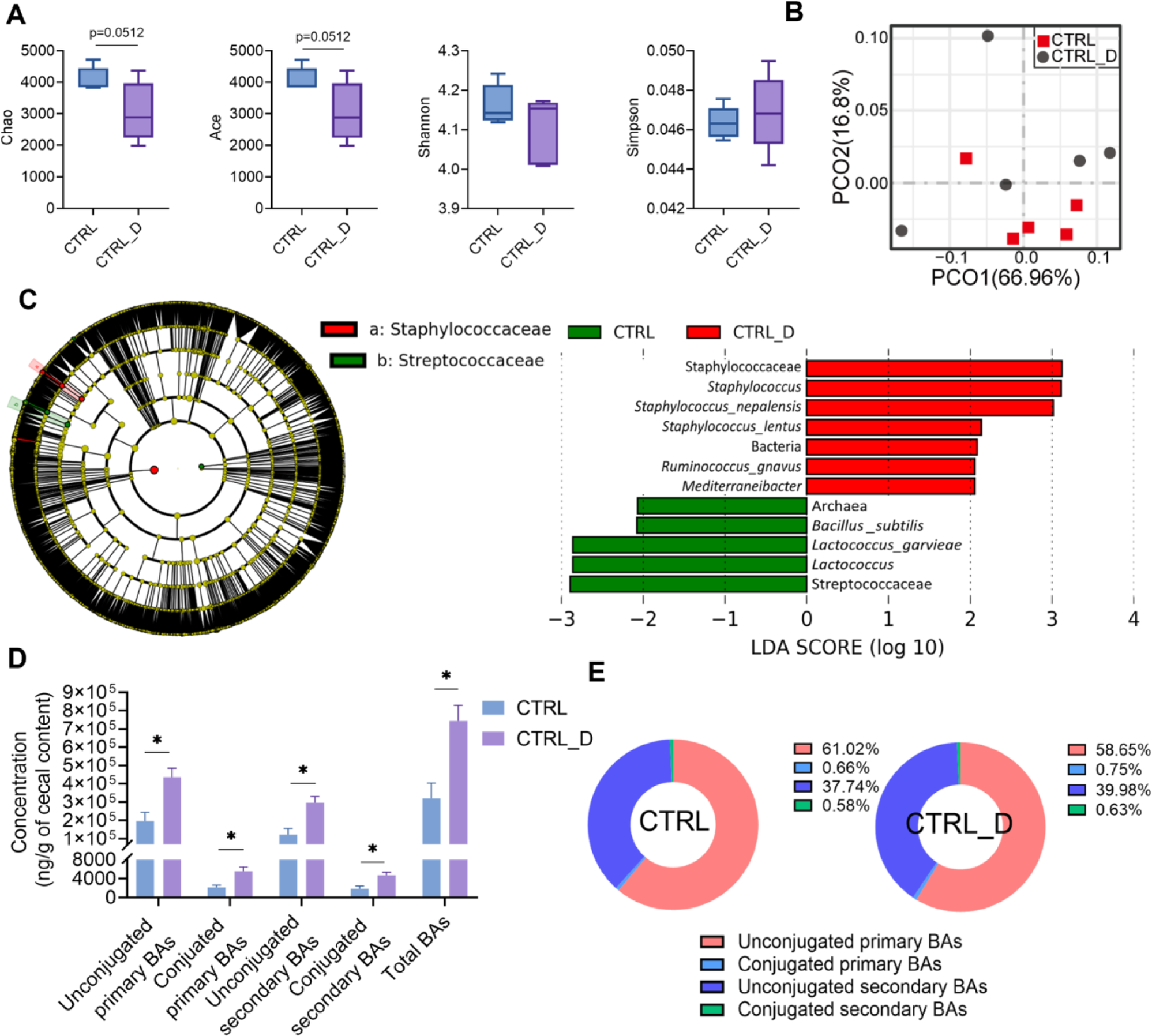
Structure and BA metabolism of gut microbiota in mice fed with the chow diet and administrated with DSV. (A) Alpha diversity. (B) PCoA (Bray-Curtis). (C) Cladogram and LDA score of LEfSe. OTUs with LDA score >2 were screened and represented the microbial biomarkers in that group. (D) Concentrations of unconjugated primary BAs, conjugated primary BAs, unconjugated secondary BAs, conjugated secondary BAs and total BAs. (E) Percentage composition of BAs. Mean values of each BA type were used to draw the pie charts. n = 5 mice per group in (A-C); n = 3-5 mice per group in (D-E). Data are represented as the mean ± SEM, two-tailed Student t-test or Mann-Whitney test,\**p* < 0.05.

Nevertheless, DSV administration was enough to alter the BA metabolism of the gut microbiota probably due to the effect of certain metabolites or structural components of DSV. The unconjugated primary BAs, conjugated primary BAs, unconjugated secondary BAs and conjugated secondary BAs all significantly increased in the CTRL_D group, which indicated enhanced host BA production and more active microbial BA metabolism (Fig. 4D). Concentrations of primary BAs including apocholic acid (ACA), chenodeoxycholic acid (CDCA), ursodeoxycholic acid (UDCA), tauro-alpha-muricholic acid (α-TMCA) and tauro-beta-muricholic acid (β-TMCA) and secondary BAs including 3-DHCA, 7-KLCA, hyocholic acid (HCA), β-HDCA, ω-TMCA and taurolithocholic acid (TLCA) remarkably increased in the CTRL_D group (Fig. S8). Our results are consistent with the study carried out by Hu *et al.*, where both the gallstone patients who had enriched Desulfovibrionales in their gut and gall-resistant mice receiving FMT from gallstone patients have increased cecal secondary BA production, which leads to increased hydrophobicity of BA pool^35^. In addition, their study shows that the expression of *Cyp7a1* is inhibited by H_2_S produced by *Desulfovibrio* species in gallstone-resistant mice when receiving extra *Desulfovibrio* ^35^, which is also consistent with the results generated in our study where hepatic *Cyp7a1* was inhibited in the CTRL_D group (Fig. 3J). However, the lower expression of *Cyp7a1* could also be explained by the negative feedback to the increased BAs in the mouse gut.

Gut barrier function was also assessed by determining the relative gene expression and IHC staining of Zo1 and Ocln of the colon tissues. Relative expression of both *Zo1* (p=0.0699) and *Ocln* was lower in the CTRL_D group though only the IHC staining of ZO1 showed significantly lower signal in the CTRL_D group (Fig. 3K-M). The elevated BA concentration together with the LPS produced by DSV were the risk factors that compromised gut barrier function and allowed translocation of toxic compounds to the liver, which induced inflammation and hepatocellular necrosis.

### SAAL diet protected the liver from injury caused by excess Desulfovibrio in the gut

To see if any dietary intervention could ameliorate the hepatic injury caused by excess *Desulfovibrio*, we compared the major nutrient contents of the chow diet and purified diet. The suggested vitamin and mineral contents for chow diet are all higher than those in the purified diet but their bioavailability is unknown since they are usually in the bound forms (Table S1). In addition, vitamins and minerals are not counted for energy. Therefore, we decided not to look at vitamins and minerals in this study. Though a few nutrients may account for the difference in gut microbiota and liver functions, we decided to focus on the amino acids because individual amino acids were added in the purified diet in our study which made their bioavailability higher and may be easier to have impacts on host health. A major difference we found between the purified and chow diet is that the content of sulfur-containing amino acids was much higher in the purified diet (Table S1), which coincides with the over-consumption of red meat in the Western style diet^36^. Therefore, to investigate whether restriction of sulfur-containing amino acids may reverse the negative effects brought by purified diet and extra DSV administration, we fed mice with the SAAL diet, which used the same premixed feed as the NRC95 standard purified diet and the two diets were isoenergetic (Fig. 5A).

**Fig. 5.**
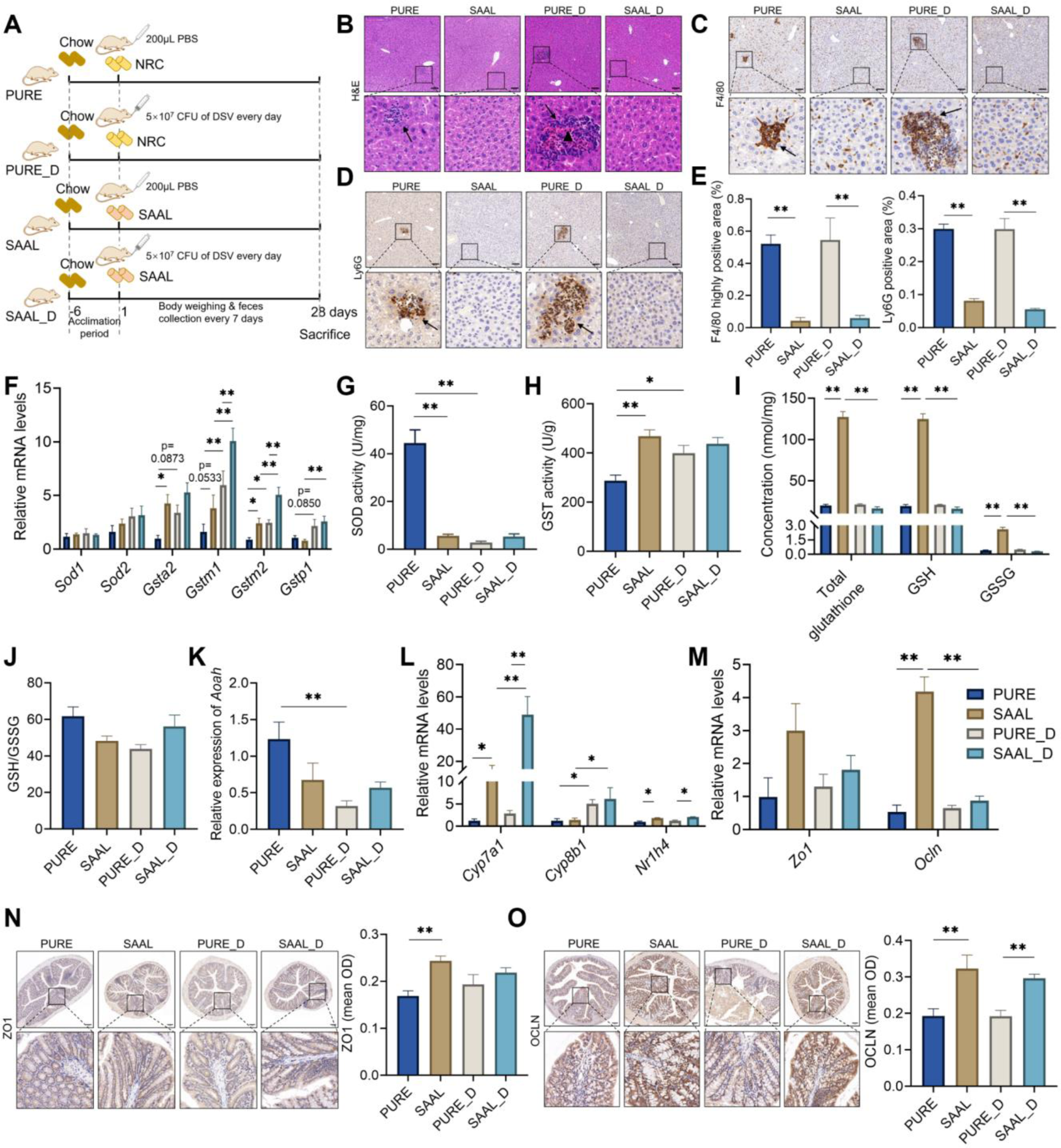
Liver and colonic changes in mice fed with the purified diet and SAAL diet with or without DSV administration. (A) Experimental design. (B) H&E staining of liver sections. (C-D) F4/80 and Ly6G IHC staining, respectively. Scale bar: 100μm. Black arrows point to the accumulation of immune cells in (B-D). Black triangle indicated hepatocellular necrosis in (B). (E) Quantification of positive areas in F4/80 and Ly6G IHC staining. (F) Relative mRNA levels of *Sod1*, *Sod*2, *Gsta2*, *Gstm1*, *Gstm2* and *Gstp1*. (G) Hepatic SOD activity. (H) Hepatic GST activity. (I) Concentrations of hepatic total glutathione, GSH and GSSG. (J) GSH/GSSG ratio. (K) Relative mRNA levels of hepatic *Aoah*. (L) Relative mRNA levels of hepatic *Cyp7a1*, *Cyp8b1* and *Nr1h4*. (M) Relative mRNA levels of *Zo1* and *Ocln* in the mouse colon. (N) IHC staining of ZO1 and quantification of mean OD of the signal. (O) IHC staining of OCLN and quantification of mean OD of the signal. Scale bar: 100μm. n = 4-6 mice per group. Data are represented as mean ± SEM, one-way ANOVA or Kruskal-Wallis test with FDR post hoc test (Two-stage step-up method of Benjamini, Krieger and Yekutieli), \**p* < 0.05, \*\**p* < 0.01.

The only difference was the reduced content of methionine (0.15%) and cystine (0%) in the SAAL diet (Table S1). Body weight of mice fed with the SAAL diet was significantly lower than that of mice fed with the purified diet no matter with or without DSV administration; in addition, the body weight of mice fed with the SAAL diet was decreasing during the experiment though the physical and mental condition of the mice looked normal (Fig. S9A). Liver/body ratio of mice in the SAAL and SAAL_D group was lower than that of mice in the PURE and PURE_D group, respectively, which was possibly due to the leaner body mass of mice fed with the SAAL diet (Fig. S9B). Daily energy intake corresponded with the body weight, in which mice fed with the purified diet had higher energy intake than mice fed with the SAAL diet did. However, DSV administration led to decreased daily energy intake in mice fed with the purified diet but increased daily energy intake in mice fed with the SAAL diet (Fig. S9C). H&E staining showed inflammatory infiltration in the PURE group but not SAAL group; DSV administration exacerbated the liver condition in which not only inflammatory infiltration but also hepatocellular necrosis was observed in the PURE_D group (Fig. 5B). Strong signals of F4/80 and Ly6G IHC staining were observed in both PURE and PURE_D group, suggesting accumulation of macrophages and neutrophils. Surprisingly, DSV administration did not lead to any histological abnormalities in mouse livers in the SAAL_D group (Fig. 5B-E).

### SAAL diet made gut microbiota in mice more resistant to DSV disruption

Alpha diversity did not show any significant difference and dimensional reduction analysis was not able to separate the microbial features between the PURE and SAAL group in metagenomic analysis (Fig. 6A-B). However, LEfSe demonstrated specific biomarkers in the PURE and SAAL group (Fig. 6C, Fig. S10A). After administration of DSV to mice, the alpha diversity still had no difference but the structure of the gut microbiota was very different between the PURE_D and SAAL_D group demonstrated by PCoA (Fig. 6A-B). The number of differential microbial biomarkers between the PURE_D and SAAL_D group increased substantially when comparing with that between the PURE and SAAL group (Fig. 6C, Fig. S10). Most of the microbial biomarkers in the SAAL_D group belonged to the order Bacteroidales, class Bacteroidia, phylum Bacteroidetes and most of the microbial biomarkers in the PURE_D group belonged to the order Clostridiales, class Clostridia, phylum Firmicutes. Therefore, Firmicutes/Bacteroidetes ratio, Clostridia/Bacteroidia ratio and Clostridiales/Bacteroidales ratio was significantly higher in the PURE_D group compared with that in the SAAL_D group (Fig. S12A-C). In addition, *Helicobacter*, which belongs to the family Helicobacteraceae, contributed to the significant increase of Proteobacteria in the PURE_D group (Fig. 6C, Fig. S10B). These findings suggest that the gut microbiota respond very differently to the DSV disturbance under the purified diet and SAAL diet.

**Fig. 6.**
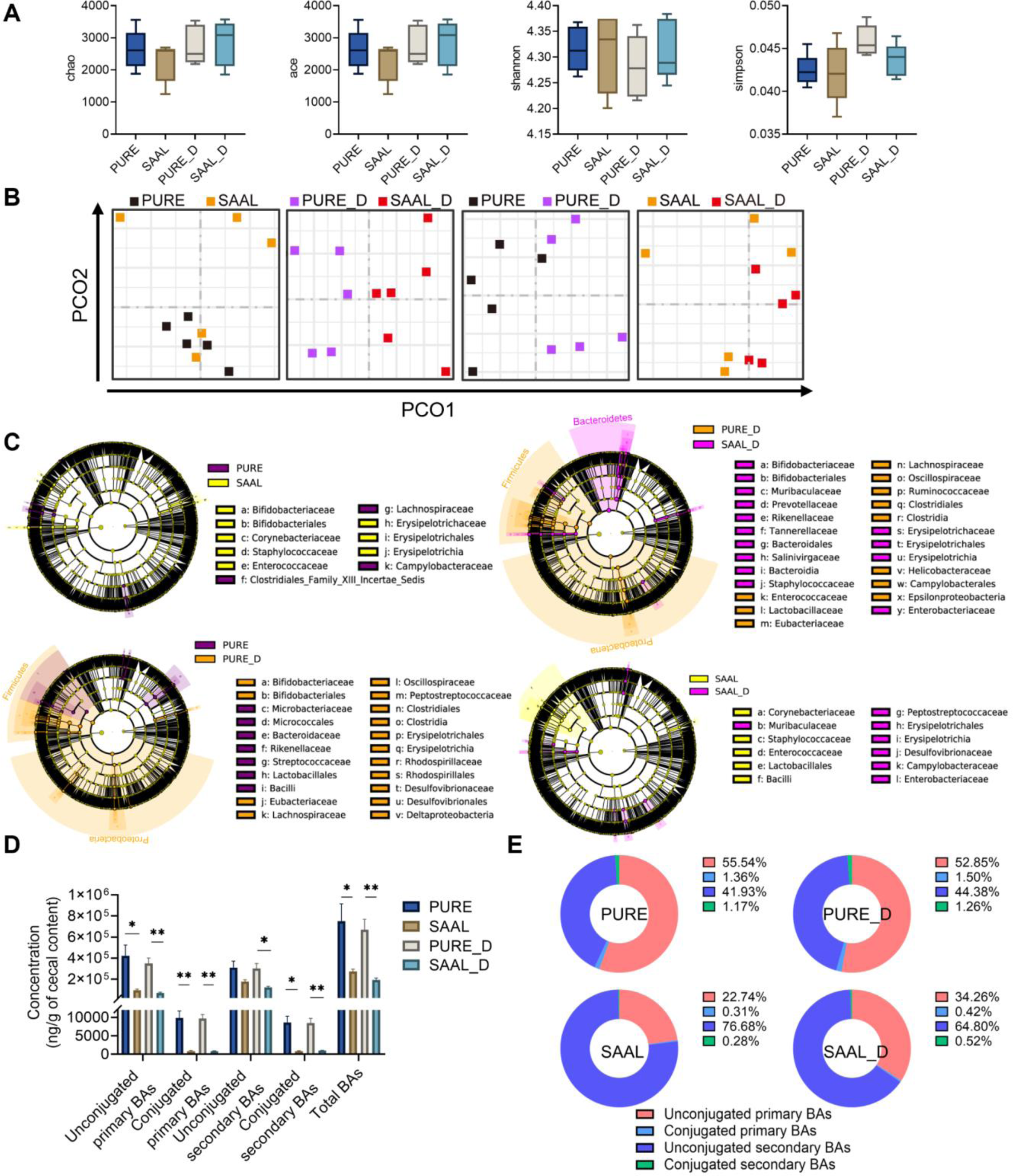
Structure and BA metabolism of gut microbiota in mice fed with the purified diet and SAAL diet with or without DSV administration. (A) Alpha diversity. (B) PCoA (Bray-Curtis). (C) Cladogram of LEfSe. OTUs with LDA score >2 were screened and represented the microbial biomarkers in that group. (D) Concentrations of unconjugated primary BAs, conjugated primary BAs, unconjugated secondary BAs, conjugated secondary BAs and total BAs. (E) Percentage composition of BAs. Mean values of each BA type were used to draw the pie charts. n = 5-6 mice per group. Data are represented as the mean ± SEM, one-way ANOVA or Kruskal-Wallis test with FDR post hoc test (Two-stage step-up method of Benjamini, Krieger and Yekutieli),\**p* < 0.05, \*\**p* < 0.01.

The composition of the gut microbiota experienced a great shift in mice fed with the purified diet after DSV administration, demonstrated by a large number of differential microbial biomarkers between PURE and PURE_D group in LEfSe (Fig. 6C, Fig. S11A). Since most of the microbial biomarkers in the PURE_D group were under the order Clostridiales, class Clostridia and phylum Firmicutes, the Firmicutes/Bacteroidetes ratio, Clostridia/Bacteroidia ratio and Clostridiales/Bacteroidales ratio got a significant increase when comparing with that in the PURE group (Fig. S12A-C). Importantly, DSV administration did increase the relative abundance of Deltaproteobacteria, Desulfovibrionales, Desulfovibrionaece and *Desulfovibrio* in the gut of mice fed with the purified diet; but interestingly, at the species level, only the relative abundance of *D. fairfieldensis* increased significantly, which suggests that the administration of extra *D. desulfuricans* may promote the growth of other *Desulfovibrio* species (Fig. S11A).

On the other hand, DSV administration did not exert such a great impact on the structure of gut microbiota in mice fed with the SAAL diet. Dimensional reduction analysis did not separate the SAAL and SAAL_D group and LEfSe showed much fewer number of microbial biomarkers (Fig. 6B-C, Fig. S11B). Similar to the case in the PURE_D group, DSV administration also led to a significant increase in the relative abundance of Desulfovibrionaece, *D. fairfieldensis* and *D. sp G11* in the SAAL_D group (Fig. S11B). Though the effect of DSV administration on gut microbial structure was different in mice fed with the purified diet and SAAL diet, it did simultaneously increase the relative abundance of Peptostreptococcaceae, Desulfovibrionaceae, Erysipelotrichales, *Helicobacter* and *Desulfovibrio* species in both groups. These results demonstrate that the purified diet created a gut environment that was easier for *Desulfovibrio* to colonize but restriction of sulfur-containing amino acids made the mouse gut more resistant to DSV disruption with less changes in the structure of the gut microbiota.

### SAAL diet reversed the elevated gut BA level due to purified diet and extra Desulfovibrio

Mouse cecal BA profile was determined. The cecal BA concentration in the SAAL group was similarly low when compared with the BA concentration in the CTRL group in our first and second experiment (Fig. 1F, Fig. 4D, Fig. 6D). However, both primary and secondary BAs in the PURE group increased significantly compared with those in the SAAL group (Fig. 6D). Though the abundance of *Desulfovibrio* which was considered to play a role in gut BA metabolism, did not decrease in mice fed with the SAAL diet, different types of BAs were still maintained in low concentrations, which indicated that other bacteria in this group may be more important in maintaining the BA homeostasis. Administration of extra DSV to mice fed with the SAAL diet did not elevate the levels of both primary and secondary BAs, which was not the case in the CTRL_D group in our second experiment (Fig. 4D, Fig. 6D). Nevertheless, DSV administration to mice fed with the purified diet also did not further increase the levels of all types of BAs (Fig. 6D, Fig. S13). This was probably due to the initially high BA level and extra DSV could not further shift the host and microbial BA metabolism. Specifically, the concentration of primary BAs including β-MCA, β-UDCA, UDCA, CDCA, MoCA, α-TMCA, β-TMCA and TCDCA and secondary BAs including IsoDCA, IsoLCA, UCA, DCA, ω-MCA, β-HDCA, LCA, HDCA, ω-TMCA, TDCA, THDCA and TLCA all remarkably increased in the PURE and PURE_D group when comparing with the SAAL and SAAL_D group, respectively (Fig. S13). Interestingly, the percentage composition of the gut BA profile in mice fed with the SAAL diet was very different from that in mice fed with the purified diet. The percentage of unconjugated secondary BAs was quite high in the SAAL and SAAL_D group (Fig. 6E). This indicated that the microbial modification of BAs was active in the mice of SAAL and SAAL_D group though the actual concentration of secondary BAs was much lower compared with that in the PURE and PURE_D group, respectively, which was due to the lower amount of conjugated primary BAs that reached to the cecum for microbial modification (Fig. 6D).

Gene expression levels of hepatic *Cyp7a1*, *Cyp8b1* and *Nr1h4* were assessed to see the effect of SAAL diet on host BA production. The expression level of hepatic *Cyp7a1* significantly increased in mice fed with the SAAL diet compared with that of mice fed with the purified diet no matter DSV was administrated or not (Fig. 5L). Higher *Cyp7a1* level indicated more active BA synthesis, which was very possibly due to compensation for the increased excretion of BAs. Though BAs excreted in feces was not determined in this study, previous research has shown that supplementation of a probiotics mix containing *Lactobacillus*, *Bifidobacterium* and *Streptococcus* species increased fecal excretion of BAs due to more active BA deconjugation that increases the hydrophobicity of the BA pool and upregulated *Cyp7a1* expression in the mouse liver to enhance hepatic neosynthesis of BAs^37^. Significantly higher abundance of *Bifidobacterium* was also found in the SAAL and SAAL_D group compared with the PURE and PURE_D group, respectively in our study and could partially explain the increased expression of *Cyp7a1* (Fig. S10, Fig. 5L). In addition, the active transformation of primary BAs into secondary BAs in mice fed with the SAAL diet (Fig. 6E) may further increase the fecal excretion of BAs since secondary BAs are usually more hydrophobic and therefore are less absorbed^26, 29, 38, 39^. The even higher *Cyp7a1* level in the SAAL_D group (Fig. 5L) may suggest more fecal excretion of BAs in this group to maintain the BA homeostasis since there was no significant elevation of cecal BA content after DSV administration to mice in the SAAL_D group (Fig. 6D). Although the expression level of *Nr1h4* in the SAAL and SAAL_D group was higher than that in the PURE and PURE_D group, respectively, which should inhibit the expression of *Cyp7a1*, the expression level of *Cyp7a1* was still higher in mice fed with the SAAL diet, which suggested that other regulations may be involved.

To further investigate the possible reason for the enhanced expression of *Cyp7a1* in SAAL_D group after DSV administration (Fig. 5L), serum cholesterol level was determined since the first step of BA synthesis is hydroxylation of cholesterol by CYP7A1, the rate-limiting enzyme. Serum LDL-C level increased remarkably in the PURE_D group while SAAL diet rescued the increased LDL-C level induced by DSV administration by enhancing the gene expression of *Cyp7a1* to expend the cholesterol for BA synthesis (Fig. 5L, Fig. S14B).

### SAAL diet protected the mouse liver from translocated toxins through enhanced antioxidant and detoxifying capability

The gene expressions of Zo1 and Ocln in the mouse colon were determined and their corresponding IHC stainings were also performed. Mice in the SAAL group possessed a better gut barrier function compared with that of mice in the PURE group, in which the expression level of *Ocln* was significantly higher and the signal of ZO1 and OCLN IHC staining was also remarkably higher (Fig. 5M-O). Higher abundance of harmful *Helicobacter* and *Campylobacter* which produce pro-inflammatory LPS and higher BA concentration especially the unconjugated secondary BAs which are more toxic damaged the gut barrier of mice in the PURE group (Fig. S10A, Fig. 6D). Translocation of these toxins through the damaged gut barrier to the liver led to liver inflammation (Fig. 5B-E). DSV administration did not make the already damaged gut barrier worse in the PURE_D group but the increase of potentially pathogenic bacteria including LPS-producing *Helicobacter* and *Desulfovibrio* species and toxin-producing *Clostridioides difficile* may increase the amount of translocated toxic compounds to the liver and therefore led to more severe hepatic injury in the PURE_D group (Fig. S11A, Fig. 5B-E).

DSV administration did disrupt the gut barrier function of mice in the SAAL_D group, in which a great decrease in the expression of *Ocln* occurred though IHC result of OCLN did not show significant difference between the SAAL and SAAL_D group (Fig. 5M-O). The BA concentration did not elevate in the SAAL_D group (Fig. 6D, Fig. S13), which means the potential damage on the gut barrier was not due to the toxic BAs. The increased abundance of potentially harmful bacteria including *Campylobacter*, *Helicobacter* and *Desulfovibrio* in the SAAL_D group, which are all gram-negative bacteria that can produce LPS and induce inflammation, were detrimental to the barrier function and may allow translocation of toxins to the liver (Fig. S11B). However, there was no any obvious damage to the mouse livers in the SAAL_D group (Fig. 5B-D).

Antioxidant and detoxifying capability was assessed to see if it was involved in the hepatoprotective effect of the SAAL diet. A tremendous decrease of SOD activity in the PURE_D group compared with the PURE group was observed, suggesting impaired antioxidant ability after DSV administration in mice fed with the purified diet (Fig. 5G). Interestingly, the SOD activity of SAAL group was much lower than that of the PURE group, probably due to the overall lower oxidative stress in mice fed with the SAAL diet and DSV administration did not impair the SOD activity in the SAAL_D group (Fig. 5G). The mRNA level of *Gsta2* and *Gstm2* was significantly higher in the SAAL group compared with the PURE group, corresponding to the higher overall GST activity in the SAAL group (Fig. 5F and H). DSV administration activated the mRNA transcription of GST genes in both PURE_D and SAAL_D group compared with that in the PURE and SAAL group, respectively, which could be a positive response to combat the elevated oxidative stress induced by extra DSV (Fig. 5F). However, only liver GST activity in the PURE_D group enhanced significantly compared with that in the PURE group (Fig. 5H). The possible reason for that GST activity did not further increase in the SAAL_D group could be that the GST activity was high enough and reached the upper limit of the enzyme activity (Fig. 5H). In addition, the mRNA levels of *Gstm1* and *Gstm2* in the SAAL_D group was remarkably higher than those in the PURE_D group, indicating even enhanced transcription of GST genes (Fig. 5F). Concentration of reduced (GSH) and oxidized form (GSSG) of glutathione and GSH/GSSG ratio were also determined. The results suggested that GSH level was much higher in the SAAL group but after DSV administration, it decreased tremendously in the SAAL_D group (Fig. 5I). Nevertheless, the GSSG level also decreased in the SAAL_D group, which made the GSH/GSSG ratio similar to that in the SAAL group (Fig. 7I-J). This means that a large amount of GSH was not converted into GSSG but very possibly devoted to the antioxidant and detoxifying activity of GSTs for conjugation with toxins, which corresponded to the enhanced transcription of GST genes in the SAAL_D group. Furthermore, the relative expression of *Aoah* greatly decreased in the PURE_D group compared with that of the PURE group, indicating compromised LPS-detoxifying ability (Fig. 5K). However, no significant difference was observed between the *Aoah* level in the SAAL and SAAL_D group (Fig. 5K).

**Fig. 7.**
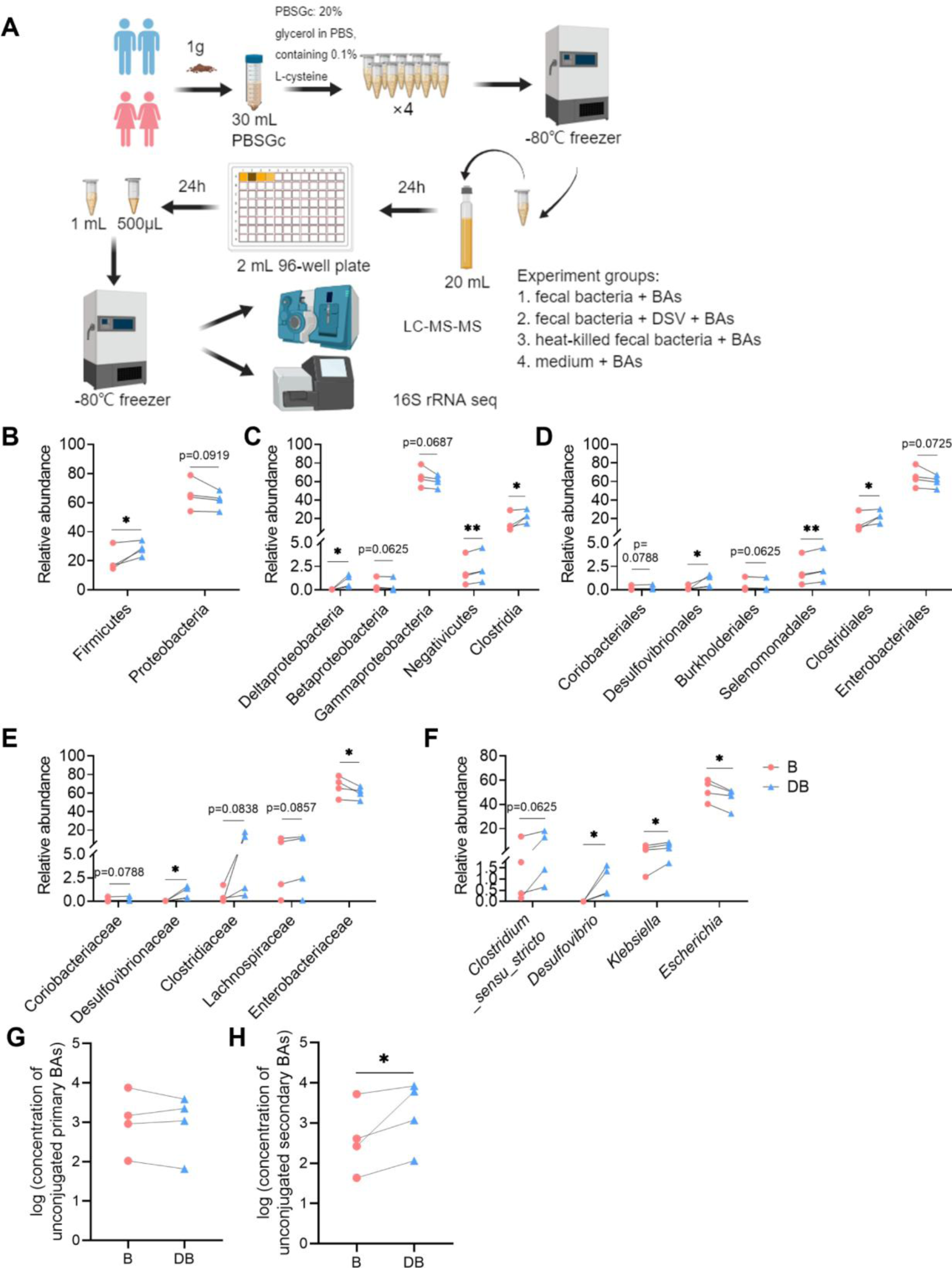
Alterations of microbial composition and BA metabolism in *ex vivo* fermentation of human fecal bacteria with primary BAs. (A) Experimental design. (B) Relative abundance of OTUs at phylum level. (C) Relative abundance of OTUs at class level. (D) Relative abundance of OTUs at order level. (E) Relative abundance of OTUs at family level. (F) Relative abundance of OTUs at genus level. (G) Log of the concentration of unconjugated primary BAs. (H) Log of the concentration of unconjugated secondary BAs. n = 4 human fecal samples per group. Data are represented as the mean ± SEM, one-tailed paired t-test or Wilcoxon matched-pairs signed rank test, \**p* < 0.05, \*\**p* < 0.01.

Therefore, the reason that the mice in the SAAL_D group could escape from the liver damage caused by the translocated toxins from the injured gut barrier after DSV administration was on account of the high antioxidant and detoxifying capability.

### Excess Desulfovibrio in human gut microbiota elevated the production of unconjugated secondary BAs

To see whether the excess DSV could also alter the human gut microbial BA metabolism, fecal samples of 4 healthy volunteers were collected and fermented with four primary BAs (Fig. 7A). 16s rRNA gene sequencing was first performed to determine if there was any change in micrbial structure after adding extra DSV in the *ex vivo* bacterial culture. Alpha diversity analysis showed increased Shannon index and decreased Simpson index, indicating increased diversity of bacteria in the DB group (Fig. S15). The relative abundance of OTUs at each taxonomic level of the two groups were compared (only OTUs with relative abundance > 0.5% were included). Increased relative abundance of Deltaproteobacteria, Desulfovibrionales, Desulfovibrionaceae and *Desulfovibrio* in the DB group was within expectation since extra DSV was added in the samples (Fig. 7C-F). The relative abundance of order Clostridiales, class Clostridia and phylum Firmicutes also increased significantly in the DB group (Fig. 7B-D). Studies have suggested that a few bacteria belonging to order Clostridiales have active BA metabolism which own both bai operons possessing activity of 7α-dehydroxylation and hydroxysteroid dehydrogenases (HSDHs) possessing activity of epimerization and oxidation^29, 40^. In addition, *Klebesiella*, a well-researched virulent bacterium that produce LPS, also increased significantly in the DB group (Fig. 7F). *Klebesiella* has been regularly reported for its association with gut diseases such as IBD^41, 42^.

The microbial BA metabolic activity including deconjugation and hydroxylation of primary BAs was determined by quantification of unconjugated primary BAs including CA and CDCA and commonly found unconjugated secondary BAs including LCA, DCA, UDCA and HDCA in the fecal bacterial culture (Fig. 7G-H, Fig. S16). When extra DSV was added in the fecal bacteria, the total concentration of four unconjugated secondary BAs increased significantly in the DB group, which indicated more active bacterial BA metabolism (Fig. 7H). The increased relative abundance of BA-metabolizing Clostridiales could partially explained the increased unconjugated secondary BA level. Last but not least, only a small amount of unconjugated primary BAs (CA only) was detected in the two negative controls which were the heat-killed bacterial culture and abiotic medium, probably due to some nonenzymatic breakdown; no unconjugated secondary BAs were detected, indicating that only live bacteria own the enzymatic activity to metabolize BAs (Fig. S17).

### DSV enhanced BA production and inhibited Cyp7a1 expression in germ-free mice

To assess if *Desulfovibrio* itself could affect host BA metabolism, 2×10^8^ CFU of DSV was administrated to germ-free mice for 14 days (Fig. 8A). The levels of primary BAs including TCA, TCDCA, TUDCA, GCA, GCDCA, GUDCA, CA, CDCA, α-MCA and β-MCA and secondary Bas including ω-MCA, DCA, LCA and HDCA were determined in the mouse cecal content. Gut BA profile of germ-free mice was quite different compared to that of conventional mice. Conjugated primary BAs including TCA, TCDCA and TUDCA were detected while GCA, GCDCA and GUDCA were not detectable in the GFC group. Only minimal concentration of unconjugated primary BAs including α-MCA and β-MCA was detected while CA, CDCA and UDCA was not detectable. In addition, the detected concentration of secondary BAs including LCA and DCA was negligible and only minimal amount of ω-MCA was detected in the GFC group (Fig. 8C). This result suggested that deconjugation of conjugated primary BAs and production of secondary BAs was almost impossible in mice without gut microbiota, which corresponds with the results of previous researches^43, 44^. DSV administration enhanced the concentration of TCA in the cecum, indirectly reflecting higher host production of BAs (Fig. 8C). Increased concentration of α-MCA and ω-MCA after DSV administration in the GFDSV group suggested that *Desulfovibrio* possessed the ability to deconjugate conjugated primary BAs and the ability to modify primary BAs to produce secondary BAs (Fig. 8C). Gene expression of hepatic *Cyp7a1* was reduced in GFDSV group, which could be the negative feedback to the enhanced concentration of TCA, which is a FXR agoist (Fig. 8B). Though the expression of *Nr1h4* did not change in the GFDSV group, the expression of small heterodimer partner *(Shp*) increased, whose expression is induced by FXR (encoded by *Nr1h4*) and subsequently inhibits the expression of *Cyp7a1*^26^. These results suggested that *Desulfovibrio* itself could affect the host BA metabolism without other gut microbes. Nevertheless, the significantly altered gut BA profile in conventional mice still relied on the effect of *Desulfovibrio* on the whole gut microbiota.

**Fig. 8.**
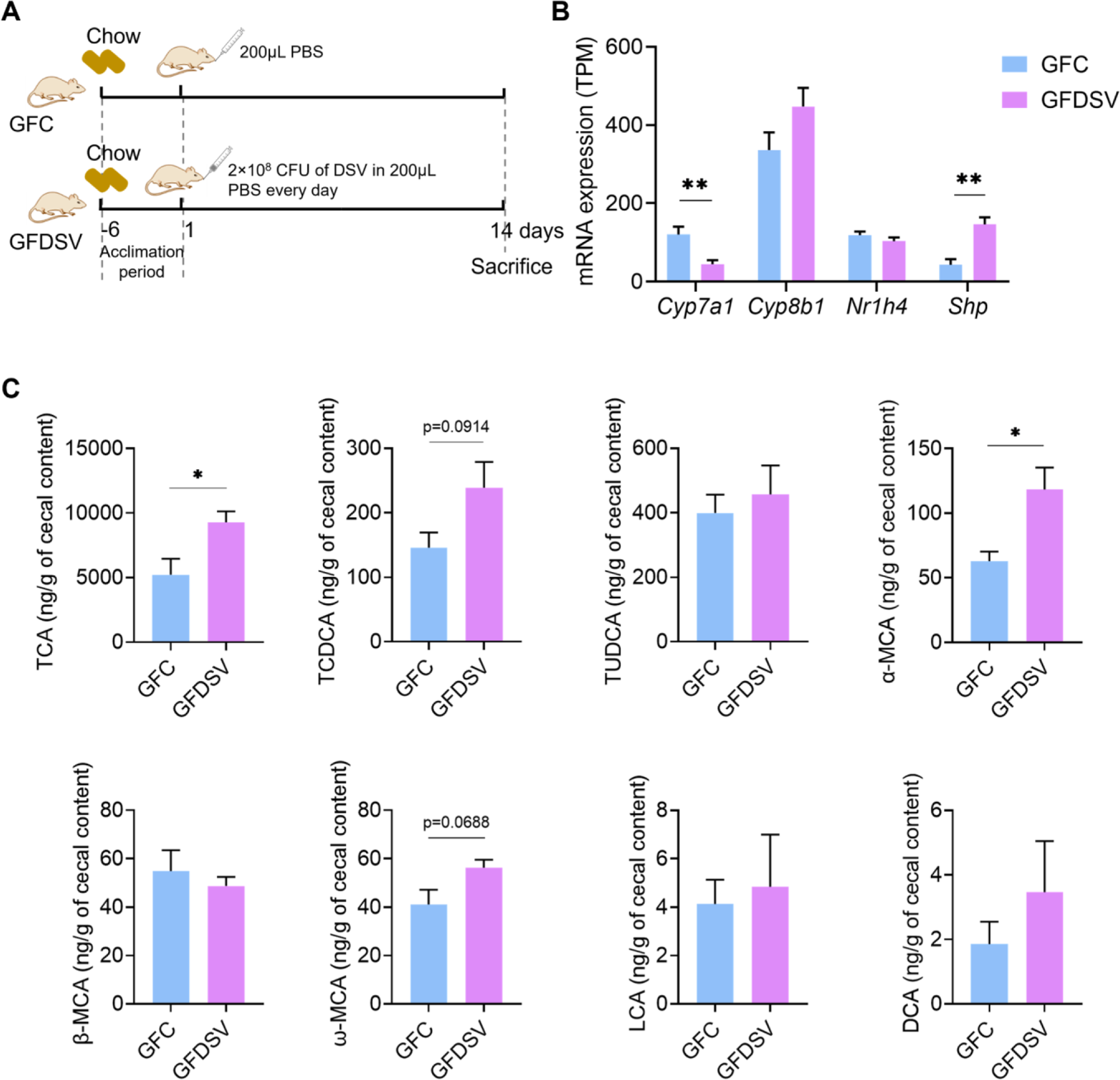
Effect of DSV on host BA synthesis and gut BA profile in germ-free mice. A. mRNA expression of *Cyp7a1*, *Cyp8b1*, *Nr1h4* and *Shp* determined by RNA-seq. B. Concentration of TCA, TCDCA, TUDCA, α-MCA, β-MCA, ω-MCA, LCA and DCA. GCA, GCDCA, GUDCA, CA, CDCA and HDCA were under detection limit in both groups. UDCA was also not shown because it was detected in the GFDSV group but not detected in the GFC group. n=4 mice per group. Data are represented as the mean ± SEM, two-tailed Student t-test, \**p* < 0.05, \*\**p* < 0.01.

## Discussion

Diet has always been the study focus for gut-liver axis since it plays an important role in maintaining structural and metabolic homeostasis of gut microbiota which largely affects the gut barrier function and determines the endotoxic and oxidative stress in the liver through translocation of pathogen-associated molecular patterns (PAMPs) including bacteria and bacterial products^1, 2, 11^. In this study, we firstly found that compared with the standard chow diet, the purified diet significantly shifted the structure of gut microbiota, enriched *Desulfovibrio*, and enhanced gut BA concentration in mice. In addition, liver inflammation and altered liver functions were also found in mice fed with the purified diet. We then demonstrated that extra DSV administration led to liver injury including inflammation and hepatocellular necrosis, impaired the liver antioxidant ability and significantly increased both primary and secondary BAs in the gut of mice fed with the chow diet. Next, our results showed that SAAL diet rescued the liver injury caused by purified diet and excess *Desulfovibrio* by enhancing detoxifying and antioxidant ability and reversed the altered gut microbial BA profile. Furthermore, we found extra *Desulfovibrio* altered the BA metabolism of bacteria in human gut through *ex vivo* fermentation of human fecal bacteria with primary BAs. Increased production of unconjugated secondary BAs and increased relative abundance of Clostridiales which contains a number of species with active BA metabolizing activity were detected in the bacterial culture after adding extra DSV in. This result correlated with what we observed in the animal experiments in which excess DSV altered the gut microbial BA metabolism and increased the production of secondary BAs. Last but not least, we also revealed that *Desulfovibrio* itself could affect the BA metabolism in germ-free mice fed with the chow diet by increasing the production of primary BAs, which in turn inhibited the expression of hepatic *Cyp7a1* as the negative feedback. Nevertheless, the effect of *Desulfovibrio* on host and gut BA metabolism was more prominent when gut microbiota were present.

When digging in the possible reasons for liver injury caused by excess *Desulfovibrio* due to diet or artificial DSV administration, the shared risk factors are increased proinflammatory LPS and toxic BAs in the gut. Increased abundance of LPS-producing Proteobacteria including *Desulfovibrio* and *Helicobacter* was observed in mice fed with the purified diet. Though DSV administration to mice fed with the chow diet did not increase the relative abundance of *Desulfovibrio* in the gut, the oxidative stress was still enhanced due to LPS produced by the administrated DSV. LPS is able to induce inflammatory signaling pathway, which damages the gut barrier and eventually leads to systemic low-grade inflammation^1, 11, 45, 46^. Previous studies have shown that LPS activates immune cells such as macrophages and neutrophils and other types of cells such as hepatic stellate cells (HSCs) to secret the proinflammatory cytokines in different liver diseases including NAFLD, NASH, ALD, liver fibrosis, liver cirrhosis and acute liver injury^1, 2, 45–52^.

BAs, as the other risk factor for liver injury, are naturally toxic and hydrophobicity of BAs decides their toxicity^26^. Secondary BAs which are usually more hydrophobic than their corresponding primary BAs may pose more noxious effects^29^. BAs help maintain the healthy gut, regulate host energy metabolism and modulate immune responses when in appropriate physiological concentrations^1, 2, 5, 26, 29^. However, excessive BAs can bring adverse effects in the gastrointestinal and hepatic diseases. Increased secondary BAs including DCA and LCA in the gut has been found in colitis and colorectal cancer models, which induces inflammatory signaling pathways, increases the oxidative damage to cellular DNA and mitochondria and activates the apoptotic pathways^53–57^. Studies have also demonstrated that excessive BAs especially the more toxic hydrophobic ones are detrimental to the liver cells^58^. Disrupted BA homeostasis has been shown in many liver diseases including cholestasis, NAFLD, NASH and hepatocellular carcinoma, in which BAs can initiate the inflammatory responses such as recruiting immune cells, inducing secretion of cytokines from Kupffer cells and chemokines from hepatocytes, directly upregulating the transcriptional factor early growth response factor 1 to initiate the inflammatory signaling, promoting transformation of antigen presenting cells from tolerogenic phenotype to immunogenic phenotype, and activating HSCs to produce pro-inflammatory and tumor-promoting mediators^2, 26, 58–67^. In our study, BA content especially the content of unconjugated secondary BAs was significantly elevated in the gut of mice fed with the purified diet no matter administrated with DSV or not and mice fed with the chow diet administrated with DSV. In short, significantly elevated BA content together with the proinflammatory LPS produced by gram-negative Proteobacteria in the mouse gut disrupted the gut barrier integrity and allowed translocation of these toxins to the liver, which led to liver injury.

Restricted intake of sulfur-containing amino acids was effective in preventing liver injury and reversed the elevated gut BA content due to extra *Desulfovibrio*. The upregulation of *Cyp7a1* was a possible mechanism for mice fed with the SAAL diet to maintain the BA homeostasis, which compensated for the increased loss of hydrophobic BAs in feces. The further increased expression of *Cyp7a1* in mice fed with the SAAL diet after DSV administration was to expend more cholesterol in order to maintain the normal serum LDL-C level which was otherwise elevated in mice fed with the purified diet after DSV administration. DSV administration also damaged the gut barrier of mice in the SAAL_D group. However, livers of mice in the SAAL_D group escaped from the damages caused by the possible translocated endotoxins from the injured gut barrier through enhanced liver antioxidant and detoxifying capability. The large reservoir of GSH together with the enhanced GST gene expressions in the liver was the major mechanism for mice fed with the SAAL diet to escape from the liver injury. GSTs play an important role in cell protection against multiple exogenous and endogenous toxic compounds, xenobiotics, products from oxidative metabolism, which prevents cells from oxidative damages through conjugation activity with GSH to reduce the toxic level of the compounds and facilitate their discharge from the cells^68, 69^. Besides classical GSH conjugation activity, GSTs also possess glutathione peroxidase activity that can reduce the organic hydroperoxides produced by lipid peroxidation and oxidative damages of DNA^68^.

Nevertheless, there are still a few limitations of this study. One of the major limitations is that the signaling pathways were not elucidated for how extra *Desulfovibrio* affects the gut microbial BA metabolism which can be further investigated through prokaryotic transcriptomic study. Another limitation is that the human fecal bacteria used in the *ex vivo* study may not represent the true microbiota in the human intestine since in fact the microbiota varied along the intestine and is very different from that in the feces^70^. However, fecal samples are more convenient to obtain and easier to comply the ethics; therefore, fecal samples are still commonly used to study the human gut microbiota. In the future, the ingestible devices designed by Shalon *et al.* that can collect fluid along the intestine can be used to carry out the studies related to human gut microbiota which can provide more accurate results^70^. Last but not least, only *D. desulfuricans* was used as a representative species of *Desulfovibrio* in our study because the previous research shows that the diversity of *Desulfovibrio* species decreases and *D. desulfuricans* becomes the most dominant one in patients with liver cirrhosis^71^. Our study also showed that DSV may promote the growth of *D. fairfieldensis*, which may be an invasive and pathogenic genus and can cause severe bacteremia^72^. *D. fairfieldensis*-derived outer membrane vesicles are found to damage the intestinal epithelial barrier and activate intrinsic inflammation^73^. The relationship between *D. desulfuricans* with other *Desulfovibrio* species are also worth investigating in the future to evaluate their different impacts on liver health.

Overall, our research suggests that dietary factors play an important role in liver health through gut-liver axis. *Desulfovibrio* was one of the key microbes that could partially explain the shifted microbial BA metabolism and hepatic injury. Although *Desulfovibrio* itself was shown to affect the BA metabolism in germ-free mice, its effect was magnified when gut microbiota was present in the conventional mice. Restricted intake of sulfur-containing amino acids rescued the hepatic injury through enhanced detoxifying and antioxidant ability and reversed the altered gut microbial BA profile caused by purified diet and excess *Desulfovibrio*. These findings were partially verified in the *ex vivo* fermentation of human fecal microbiota with primary BAs, in which extra *Desulfovibrio* enhanced production of secondary BAs. SAAL diet may become an effective dietary intervention in the future for hepatic injury associated with increased abundance of *Desulfovibrio* in the gut.

## Author Contributions

L. Zhou and Y. Geng and Z. Xu conceived and designed the study.

L. Zhou, G. Lu, Y. Nie and Y. Ren performed the experiments.

L. Zhou performed the data analysis and bioinformatics analysis.

L. Zhou wrote the manuscript. Y. Geng, Z. Xu and J. Shi revised the manuscript.

Y. Geng and Y. Xue supervised the study and acquired the funding for it.

## Data availability

All the data are available upon request. The raw data of 16s rRNA gene sequencing generated in the first animal experiment and *ex vivo* experiment of fermentation of human fecal bacteria with bile acids in this study have been deposited at National Center for Biotechnology Information (BioProject ID: PRJNA1066713 and PRJNA1067359, respectively). The raw data of metagenomic sequencing in the second and third animal experiment have been deposited at National Center for Biotechnology Information (BioProject ID: PRJNA1068753). RNA-seq data of mouse livers in the first and fourth animal experiment have been uploaded to National Center for Biotechnology Information (BioProject ID: PRJNA1067360 and PRJNA600663, respectively).

## References

1. Tilg, H., Adolph, T.E. & Trauner, M. Gut-liver axis: Pathophysiological concepts and clinical implications. Cell Metab 34, 1700–1718 (2022).

2. Tripathi, A. et al. The gut-liver axis and the intersection with the microbiome. Nat Rev Gastroenterol Hepatol 15, 397–411 (2018).

3. Albillos, A., de Gottardi, A. & Rescigno, M. The gut-liver axis in liver disease: Pathophysiological basis for therapy. J Hepatol 72, 558–577 (2020).

4. Wahlström, A., Sayin, Sama I., Marschall, H.-U. & Bäckhed, F. Intestinal Crosstalk between Bile Acids and Microbiota and Its Impact on Host Metabolism. Cell Metabolism 24, 41–50 (2016).

5. Collins, S.L., Stine, J.G., Bisanz, J.E., Okafor, C.D. & Patterson, A.D. Bile acids and the gut microbiota: metabolic interactions and impacts on disease. Nat Rev Microbiol 21, 236–247 (2023).

6. Tilg, H., Moschen, A.R. & Kaneider, N.C. Pathways of liver injury in alcoholic liver disease. J Hepatol 55, 1159–1161 (2011).

7. Tilg, H., Zmora, N., Adolph, T.E. & Elinav, E. The intestinal microbiota fuelling metabolic inflammation. Nat Rev Immunol 20, 40–54 (2020).

8. Trebicka, J., Macnaughtan, J., Schnabl, B., Shawcross, D.L. & Bajaj, J.S. The microbiota in cirrhosis and its role in hepatic decompensation. J Hepatol 75 Suppl 1, S67–S81 (2021).

9. Carpino, G. et al. Increased Liver Localization of Lipopolysaccharides in Human and Experimental NAFLD. Hepatology 72, 470–485 (2020).

10. Hoyles, L. et al. Molecular phenomics and metagenomics of hepatic steatosis in non-diabetic obese women. Nat Med 24, 1070–1080 (2018).

11. Fan, Y. & Pedersen, O. Gut microbiota in human metabolic health and disease. Nat Rev Microbiol 19, 55–71 (2021).

12. Cani, P.D. & Van Hul, M. Mediterranean diet, gut microbiota and health: when age and calories do not add up! Gut 69, 1167–1168 (2020).

13. Bárcena, C. et al. Methionine Restriction Extends Lifespan in Progeroid Mice and Alters Lipid and Bile Acid Metabolism. Cell Rep 24, 2392–2403 (2018).

14. Green, C.L., Lamming, D.W. & Fontana, L. Molecular mechanisms of dietary restriction promoting health and longevity. Nat Rev Mol Cell Biol 23, 56–73 (2022).

15. Miller, R.A. et al. Methionine-deficient diet extends mouse lifespan, slows immune and lens aging, alters glucose, T4, IGF-I and insulin levels, and increases hepatocyte MIF levels and stress resistance. Aging Cell 4, 119–125 (2005).

16. Ren, B. et al. Methionine Restriction Improves Gut Barrier Function by Reshaping Diurnal Rhythms of Inflammation-Related Microbes in Aged Mice. Front Nutr 8, 746592 (2021).

17. National Research Council (US) Subcommittee on Laboratory Animal Nutrition Nutrient Requirements of Laboratory Animals, Edn. Fourth Revised Edition. (National Academies Press (US), Washington (DC); 1995).

18. Daniel, N. et al. Dietary fat and low fiber in purified diets differently impact the gut-liver axis to promote obesity-linked metabolic impairments. Am J Physiol Gastrointest Liver Physiol 320, G1014–G1033 (2021).

19. Hou, Y. et al. A diet-microbial metabolism feedforward loop modulates intestinal stem cell renewal in the stressed gut. Nat Commun 12, 271 (2021).

20. Goto, H. et al. Effects of fructo-oligosaccharide on DSS-induced colitis differ in mice fed nonpurified and purified diets. J Nutr 140, 2121–2127 (2010).

21. Boussenna, A. et al. Impact of basal diet on dextran sodium sulphate (DSS)-induced colitis in rats. Eur J Nutr 54, 1217–1227 (2015).

22. Koleva, P., Ketabi, A., Valcheva, R., Ganzle, M.G. & Dieleman, L.A. Chemically defined diet alters the protective properties of fructo-oligosaccharides and isomalto-oligosaccharides in HLA-B27 transgenic rats. PLoS One 9, e111717 (2014).

23. Miles, J.P. et al. Supplementation of Low- and High-fat Diets with Fermentable Fiber Exacerbates Severity of DSS-induced Acute Colitis. Inflamm Bowel Dis 23, 1133–1143 (2017).

24. Gonzalez-Blazquez, R. et al. Relevance of control diet choice in metabolic studies: impact in glucose homeostasis and vascular function. Sci Rep 10, 2902 (2020).

25. Xie, G. et al. Distinctly altered gut microbiota in the progression of liver disease. Oncotarget 7, 19355–19366 (2016).

26. Fuchs, C.D. & Trauner, M. Role of bile acids and their receptors in gastrointestinal and hepatic pathophysiology. Nature Reviews Gastroenterology & Hepatology 19, 432–450 (2022).

27. Yuan, L. & Kaplowitz, N. Glutathione in liver diseases and hepatotoxicity. Mol Aspects Med 30, 29–41 (2009).

28. Aggarwal, S., Dimitropoulou, C., Lu, Q., Black, S.M. & Sharma, S. Glutathione supplementation attenuates lipopolysaccharide-induced mitochondrial dysfunction and apoptosis in a mouse model of acute lung injury. Front Physiol 3 (2012).

29. Cai, J., Sun, L. & Gonzalez, F.J. Gut microbiota-derived bile acids in intestinal immunity, inflammation, and tumorigenesis. Cell Host Microbe 30, 289–300 (2022).

30. Bryant, C.E., Spring, D.R., Gangloff, M. & Gay, N.J. The molecular basis of the host response to lipopolysaccharide. Nat Rev Microbiol 8, 8–14 (2010).

31. Zhang, X. et al. Dietary cholesterol drives fatty liver-associated liver cancer by modulating gut microbiota and metabolites. Gut 70, 761–774 (2021).

32. Goldstein, E.J., Citron, D.M., Peraino, V.A. & Cross, S.A. Desulfovibrio desulfuricans bacteremia and review of human Desulfovibrio infections. J Clin Microbiol 41, 2752–2754 (2003).

33. Feng, J. et al. A host lipase prevents lipopolysaccharide-induced foam cell formation. iScience 24, 103004 (2021).

34. Han, Y.H. et al. Enterically derived high-density lipoprotein restrains liver injury through the portal vein. Science 373 (2021).

35. Hu, H. et al. Gut microbiota promotes cholesterol gallstone formation by modulating bile acid composition and biliary cholesterol secretion. Nat Commun 13, 252 (2022).

36. Strate, L.L. et al. Western Dietary Pattern Increases, and Prudent Dietary Pattern Decreases, Risk of Incident Diverticulitis in a Prospective Cohort Study. Gastroenterology 152, 1023–1030 e1022 (2017).

37. Degirolamo, C., Rainaldi, S., Bovenga, F., Murzilli, S. & Moschetta, A. Microbiota modification with probiotics induces hepatic bile acid synthesis via downregulation of the Fxr-Fgf15 axis in mice. Cell Rep 7, 12–18 (2014).

38. Chen, M.L., Takeda, K. & Sundrud, M.S. Emerging roles of bile acids in mucosal immunity and inflammation. Mucosal Immunology 12, 851–861 (2019).

39. Pushpass, R.-A.G., Alzoufairi, S., Jackson, K.G. & Lovegrove, J.A. Circulating bile acids as a link between the gut microbiota and cardiovascular health: impact of prebiotics, probiotics and polyphenol-rich foods. Nutrition Research Reviews 35, 161–180 (2022).

40. Kim, K.H., et al. Identification and Characterization of Major Bile Acid 7alpha-Dehydroxylating Bacteria in the Human Gut. mSystems 7, e0045522 (2022).

41. Read, E., Curtis, M.A. & Neves, J.F. The role of oral bacteria in inflammatory bowel disease. Nature Reviews Gastroenterology & Hepatology 18, 731–742 (2021).

42. Zhang, Q.J. et al. Kebsiella pneumoniae Induces Inflammatory Bowel Disease Through Caspase-11-Mediated IL18 in the Gut Epithelial Cells. Cell Mol Gastroenter 15, 613–632 (2023).

43. Claus, S.P. et al. Systemic multicompartmental effects of the gut microbiome on mouse metabolic phenotypes. Mol Syst Biol 4, 219 (2008).

44. Swann, J.R. et al. Systemic gut microbial modulation of bile acid metabolism in host tissue compartments. Proc Natl Acad Sci U S A 108 Suppl 1, 4523–4530 (2011).

45. Cani, P.D. et al. Metabolic endotoxemia initiates obesity and insulin resistance. Diabetes 56, 1761–1772 (2007).

46. Cani, P.D. et al. Changes in gut microbiota control metabolic endotoxemia-induced inflammation in high-fat diet-induced obesity and diabetes in mice. Diabetes 57, 1470–1481 (2008).

47. Chen, S.N. et al. Deletion of TLR4 attenuates lipopolysaccharide-induced acute liver injury by inhibiting inflammation and apoptosis. Acta Pharmacol Sin 42, 1610–1619 (2021).

48. Sookoian, S. et al. Intrahepatic bacterial metataxonomic signature in non-alcoholic fatty liver disease. Gut 69, 1483–1491 (2020).

49. Fukui, H., Brauner, B., Bode, J.C. & Bode, C. Plasma endotoxin concentrations in patients with alcoholic and non-alcoholic liver disease: reevaluation with an improved chromogenic assay. J Hepatol 12, 162–169 (1991).

50. Munoz, L. et al. Intestinal Immune Dysregulation Driven by Dysbiosis Promotes Barrier Disruption and Bacterial Translocation in Rats With Cirrhosis. Hepatology 70, 925–938 (2019).

51. Sorribas, M. et al. FXR modulates the gut-vascular barrier by regulating the entry sites for bacterial translocation in experimental cirrhosis. J Hepatol 71, 1126–1140 (2019).

52. Gandhi, C.R. Pro- and Anti-fibrogenic Functions of Gram-Negative Bacterial Lipopolysaccharide in the Liver. Front Med (Lausanne) 7, 130 (2020).

53. Xu, M.Q. et al. Deoxycholic Acid-Induced Gut Dysbiosis Disrupts Bile Acid Enterohepatic Circulation and Promotes Intestinal Inflammation. Digest Dis Sci 66, 568–576 (2021).

54. Liu, L. et al. Deoxycholic acid disrupts the intestinal mucosal barrier and promotes intestinal tumorigenesis. Food Funct 9, 5588–5597 (2018).

55. Jia, W., Xie, G. & Jia, W. Bile acid-microbiota crosstalk in gastrointestinal inflammation and carcinogenesis. Nat Rev Gastroenterol Hepatol 15, 111–128 (2018).

56. Gadaleta, R.M., Garcia-Irigoyen, O. & Moschetta, A. Bile acids and colon cancer: Is FXR the solution of the conundrum? Molecular Aspects of Medicine 56, 66–74 (2017).

57. Payne, C.M., Bernstein, C., Dvorak, K. & Bernstein, H. Hydrophobic bile acids, genomic instability, Darwinian selection, and colon carcinogenesis. Clin Exp Gastroenterol 1, 19–47 (2008).

58. Woolbright, B.L., McGill, M.R., Yan, H. & Jaeschke, H. Bile Acid-Induced Toxicity in HepaRG Cells Recapitulates the Response in Primary Human Hepatocytes. Basic Clin Pharmacol Toxicol 118, 160–167 (2016).

59. Woolbright, B.L. et al. Bile acid-induced necrosis in primary human hepatocytes and in patients with obstructive cholestasis. Toxicol Appl Pharmacol 283, 168–177 (2015).

60. Song, P., Zhang, Y. & Klaassen, C.D. Dose-response of five bile acids on serum and liver bile Acid concentrations and hepatotoxicty in mice. Toxicol Sci 123, 359–367 (2011).

61. Miyake, J.H., Wang, S.L. & Davis, R.A. Bile acid induction of cytokine expression by macrophages correlates with repression of hepatic cholesterol 7alpha-hydroxylase. J Biol Chem 275, 21805–21808 (2000).

62. Sato, K. et al. Pathogenesis of Kupffer Cells in Cholestatic Liver Injury. Am J Pathol 186, 2238–2247 (2016).

63. Allen, K., Jaeschke, H. & Copple, B.L. Bile acids induce inflammatory genes in hepatocytes: a novel mechanism of inflammation during obstructive cholestasis. Am J Pathol 178, 175–186 (2011).

64. Rahman, A.H. & Aloman, C. Dendritic cells and liver fibrosis. Biochim Biophys Acta 1832, 998–1004 (2013).

65. Carambia, A. et al. TGF-beta-dependent induction of CD4(+)CD25(+)Foxp3(+) Tregs by liver sinusoidal endothelial cells. J Hepatol 61, 594–599 (2014).

66. Yoshimoto, S. et al. Obesity-induced gut microbial metabolite promotes liver cancer through senescence secretome. Nature 499, 97–101 (2013).

67. Friedman, S.L. Hepatic stellate cells: protean, multifunctional, and enigmatic cells of the liver. Physiol Rev 88, 125–172 (2008).

68. Allocati, N., Masulli, M., Di Ilio, C. & Federici, L. Glutathione transferases: substrates, inihibitors and pro-drugs in cancer and neurodegenerative diseases. Oncogenesis 7, 8 (2018).

69. Dasari, S., Ganjayi, M.S., Yellanurkonda, P., Basha, S. & Meriga, B. Role of glutathione S-transferases in detoxification of a polycyclic aromatic hydrocarbon, methylcholanthrene. Chem Biol Interact 294, 81–90 (2018).

70. Shalon, D. et al. Profiling the human intestinal environment under physiological conditions. Nature 617, 581–591 (2023).

71. Lu, G. et al. Diversity and Comparison of Intestinal Desulfovibrio in Patients with Liver Cirrhosis and Healthy People. Microorganisms 11 (2023).

72. Pimentel Jason, D. & Chan Raymond, C. Desulfovibrio fairfieldensis Bacteremia Associated with Choledocholithiasis and Endoscopic Retrograde Cholangiopancreatography. Journal of Clinical Microbiology 45, 2747–2750 (2007).

73. Nie, Y. et al. Desulfovibrio fairfieldensis-Derived Outer Membrane Vesicles Damage Epithelial Barrier and Induce Inflammation and Pyroptosis in Macrophages. Cells 12 (2022).

