## Supplementary information for "Restricted intake of sulfur-containing amino acids reversed the hepatic injury induced by excess *Desulfovibrio* through gut-liver axis"

### Supplementary methods

#### *Sample collection*

Mouse fecal samples were collected every 7 days. Mice were sacrificed after 28 days for the first three animal experiments or 14 days for the last animal experiment. Blood was collected from mouse orbits and serum was isolated by centrifugation at 1,500 × g, 4 °C for 15 minutes. Liver and cecal samples were collected and immediately put into liquid nitrogen. All samples were kept at -80 °C for later experiments.

#### *Physiological parameters*

Mouse body weight was measured every 7 days. Daily energy intake was determined by weighing the given and left feed every day. Mouse liver weight was measured after sacrifice. Serum total cholesterol and LDL-C were measured using an automatic serum biochemical analyzer (ZY310, Shanghai Kehua Bio-Engineering co., Ltd., Shanghai, China).

#### *Histological examination*

Liver tissues were fixed in 4% paraformaldehyde and embedded in paraffin. Liver section were stained with hematoxylin and eosin (H&E). Immunohistochemical (IHC) staining of F4/80 and Ly6G were also performed for liver sections. IHC staining of Zona occludens 1 (ZO1) and Occludin (OCLN) were performed for colon sections. All experiment procedures can be referred to the standard protocols on the official website of Wuhan Servicebio Technology Co., Ltd. (Wuhan, China). Images of stained liver sections were obtained by Pannoramic MIDI II Digital Slide Scanner (3D HISTECH Ltd., Budapest, Hungry).

#### *16s rRNA gene sequencing*

200 mg of cecal content of each mouse was sent to BGI Genomics Co., Ltd. (Wuhan, China) for 16s rRNA gene sequencing. Briefly, DNA was extracted and the V3-V4 region of 16s rDNA was amplified. PCR products were purified using Agencourt AMPure XP Beads and screened for quality using Agilent 2100 Bioanalyzer. Qualified DNA sequence library was constructed and sequenced by Illumina-HiSeq 2500 platform (Illumina, San Diego, CA). Clean reads were spliced into Tags using FLASH (v.1.2.11) and Tags were clustered into OTUs using USEARCH (v.7.0.1090). Sequence of each OTU was compared with the RDP database (Release11.5 20160930) using RDP classifier (v1.9.1) software for species annotation. Following analyses included alpha diversity using Mothur software (v.1.31.2), beta-diversity using QIIME software (v.1.80), linear discriminant analysis of effect sizes (LEfSe) using lefser (v.1.12.1) in R (v.4.2.0) and function prediction using PICRUST2 (V2.3.0-b) were conducted.

#### *Metagenomic sequencing*

Bacterial DNA was extracted from the mouse cecal content and metagenomic sequencing was performed by BGI Genomics Co., Ltd. (Wuhan, China) using DNBSEQ platform. Briefly, low-quality data were filtered by software SOAPnuke (v1.5.0) and host sequences were filtered using Bowtie2 (v2.2.5). Filtered clean data were assembled using MEGAHIT (v1.1.3) and fragments under 300 bp were discarded. Kraken2 (2.1.2) was used to perform the species annotation for metagenomic data to get the abundance of species against Nt database (202011). Following taxonomy analyses included alpha diversity, beta-diversity, LEfSe and so on were conducted using R packages (v.4.2.0).

***Metabolomic study***

H650 targeted metabolomic study of mouse cecal content were performed by Shanghai Applied Protein Technology Co., Ltd. (Shanghai, China). Briefly, H650 targeted metabolomic study is consist of 650 widely studied metabolites including amino acids and derivatives, lipids and derivatives, pyridines, carbohydrates and derivatives, benzene ring-containing compounds, organic acids and derivatives, bile acids, indoles and derivatives, amines and so on.

*Metabolite extraction*

To extract the metabolites, 800 μL of cold methanol / acetonitrile / water (2:2:1, v/v) extraction solvent was added to 100 mg of cecal sample following the addition of stable-isotope internal standards. Then the samples were under vigorous shaking for 2 min at 4 °C and incubated on ice for 20 minutes. Samples were then centrifuged at 14,000 × g for 20 minutes at 4 °C. The supernatant was collected and flowed through a 96-well protein precipitation plate. The elution was collected and dried in a vacuum centrifuge at 4 °C. The samples were re-dissolved in 100 μL acetonitrile/water (1:1, v/v) solvent and centrifuged at 14,000 × g at 4 °C for 15 min, and the supernatant was ready for LC-MS analysis.

*Settings for UPLC and mass spectrometry*

Analyses were performed using an UHPLC (1290 Infinity LC, Agilent Technologies) coupled to a QTRAP 6500+ (AB Sciex LLC, MA, USA). The analytes were separated on HILIC (Waters UPLC BEH Amide column, 2.1 mm × 100 mm, 1.7 μm) and C18 columns (Waters UPLC BEH C18 column, 2.1 mm × 100 mm, 1.7 μm). The column temperature was set at 35 °C and the injection volume was 2 μL for HILIC separation. Mobile phase A was 90% $H_2O$ + 2 mM ammonium formate + 10% acetonitrile and mobile phase B was 0.4% formic acid in methanol with a gradient (85% B at 0-1 min,

80% B at 3-4 min, 70% B at 6 min, 50% B at 10-15.5 min, 85% B at 15.6 -23 min). Flow rate was 300 μL/min. For RPLC separation using C18 column, the column temperature was set at 40 °C and the injection volume was 2 μL. Mobile phase A was 5 mM ammonium acetate and 0.2% $NH_3 \cdot H_2O$ in water and mobile phase B was 99.5% acetonitrile + 0.5% $NH_3 \cdot H_2O$ with a gradient (5% B at 0 min, 60% B at 5 min, 100% B at 11-13 min, 5% B at 13.1-16 min). Flow rate was 400 μL/min. The samples were placed at 4 °C during the whole process. 6500+ QTRAP (AB Sciex LLC, MA, USA) was used to perform the following mass spectrometry in positive and negative switch mode. The settings of both modes were as follows: Source temperature: 580 °C; Ion Source Gas 1 (GS1): 45; Ion Source Gas 2 (GS2): 60; Curtain Gas (CUR): 35; Ion Spray Voltage (IS): +4500 V for positive mode and -4500 V for negative mode.

The QCs were processed together with the biological samples. Metabolites in QCs with coefficient of variation (CV) less than 30 % were denoted as reproducible measurements. MRM data was analyzed by Analyst software (v.1.7) to generate the quantitative data of each metabolite detected. After sum-normalization, the processed data were uploaded to SIMCA-P (version 14.1, Umetrics, Umea, Sweden), where it was subjected to multivariate data analysis, including Pareto-scaled principal component analysis (PCA) and orthogonal partial least-squares discriminant analysis (OPLS-DA). The 7-fold cross-validation and response permutation testing were used to evaluate the robustness of the model. The variable importance in the projection (VIP) value of each variable in the OPLS-DA model was calculated to indicate its contribution to the classification.

***Transcriptomic study***

Liver samples were prepared and sent to BGI Genomics Co., Ltd. (Wuhan, China) for RNA-sequencing (RNA-seq) using DNBSEQ platform. Briefly, RNA was extracted from liver samples using TRIzol-chloroform method. RNA samples were denatured to expose their secondary structure and then mRNA was enriched using oligo (dT)-attached magnetic beads. mRNA was fragmented and double-stranded cDNA was synthesized. Double-stranded cDNA was then subject to end repair by adding a single A' nucleotide to the 3' ends of the blunt fragments. At the same time adaptors were also ligated to cDNA. Single-stranded PCR products were generated by denaturation and circular DNA was produced using suitable circularization program. Single-stranded circular DNA molecules were replicated via rolling-circle amplification, and a DNA nanoball (DNB) which contained multiple copies of DNA was generated. Sufficient quality of DNBs were then loaded into patterned nanoarrays using high-density DNA nanochip technique and sequenced through combinatorial Probe-Anchor Synthesis (cPAS) technique.

The sequencing data was filtered with SOAPnuke (v1.5.6) by removing unqualified reads. Clean reads were mapped to reference genome (GCF_000001635.26_GRCm38.p6) using HISAT2 (v2.1.0) and then aligned to reference gene sets using Botwie2 (v2.3.4.3). Expression levels of genes were calculated by RSEM (v1.3.1). Differentially expressed genes were analyzed using DESeq2 (v1.4.5) with q value $\leq$ 0.05. KEGG (Release 108.0) enrichment analysis was performed for annotated genes with $\log2|FC| > 0.58$ and q value $\leq$ 0.05 using Phyper package in R (v4.3.0) based on hypergeometric test. Gene Set Enrichment Analysis (GSEA) was also performed against KEGG database for all annotated genes to determine whether a pre-defined set of genes exhibits statistical difference between two groups[1]. Statistical significance was represented by $p < 0.05$ and FDR < 0.25 for GSEA.

***Gene expressions determined by qRT-PCR***

1.5 mL TRIzol was added to 100 mg of liver samples or colon samples and homogenized at 4 °C. 300 μL chloroform was added into the samples and vortexed for 20 seconds. The samples were centrifuged at 12,000 × g, 4 °C for 15 minutes after settling at room temperature for 10 minutes. The transparent top layer was transferred into a new tube and equal volume of isopropanol was added. The samples were then gently mixed, settled at room temperature for 10 minutes and centrifuged at 12,000 × g, 4 °C for 10 minutes. The supernatant was discarded and 1.5 mL 75% ethanol (v/v) was added to resuspend the pallets. The samples were then centrifuged at 7,500 × g, 4 °C for 5 minutes and the supernatant was discarded. Short centrifugation was performed and the extra ethanol was removed by pipetting. After drying at room temperature for 5 minutes, 100μL DEPC water was added to dissolve the pallets.

cDNA was generated using 5X All-in-One MasterMix (with AccurRT Genomic DNA Removal Kit) kit (Applied Biological Materials (China) Inc., Nanjing, China). Quantitative real-time PCR (qRT-PCR) using SYBR$^{TM}$ Select Master Mix for CFX (Thermo Fisher Scientific Inc., Shanghai, China) was conducted to quantify the relative expression of genes. The reaction steps were as follows: 50 °C, 2 min, 95 °C, 2 min, 1 cycle, 95 °C, 15 s, 60 °C, 1 min, 39 cycles, with one fluorescence detection per cycle; after the reaction was completed, the melting curve was plotted by increasing the temperature from 65 °C to 95 °C at a rate of 0.5 °C every 5 seconds. Primer sequences can be found in Table S2.

***Evaluation of hepatic oxidative stress***

Reduced glutathione (GSH) and oxidized glutathione (GSSG) content were determined according to the standard protocol of GSH and GSSG Assay Kit (S0053) purchased from Beyotime

Biotechnology Inc. (Shanghai, China). Superoxide dismutase (SOD) activity was determined according to the standard protocol of Total Superoxide Dismutase Assay Kit (S0109) purchased from Beyotime Biotechnology Inc. (Shanghai, China). Glutathione S-transferase (GST) activity was determined according to the standard protocol of GST Activity Assay Kit (BC0350) purchased from Beijing Solarbio Science & Technology Co., Ltd. (Beijing, China).

#### *Ex vivo fermentation of human fecal bacteria with bile acids*

To investigate if the result obtained from the animal experiments in which excess *Desulfovibrio* changed gut microbial bile acid (BA) metabolism, could be transferred to human gut microbes, feces of 2 healthy male and 2 female volunteers were collected. Volunteers were between 23-30 years old without any metabolic diseases and did not take any antibiotics during past three months. Written informed consent was obtained from each volunteer included in the study after being informed of the purpose of the sample collection and the study protocol conforms to the ethical guidelines of the 1975 Declaration of Helsinki as reflected in a priori approval by the Human Research Committee of Jiangnan University (Ethics approval no. JNU20220310IRB47). The pretreatment of the fecal samples and *ex vivo* culturing method followed the methods in Javdan *et al.*'s paper[2]. Briefly, 1 gram of each fecal sample was mixed with 15 mL sterile pre-reduced phosphate buffer (PBS) with 0.1% L-cysteine and 15 mL 40% glycerol. After vigorous vortex, the homogenate was left still for 5 minutes to allow the insoluble particles to settle to the bottom. Several aliquots of 1 mL of the supernatant were collected and stored at -80 °C for later use. The whole process was performed in an anaerobic chamber (90% $N_2$, 5% $CO_2$ and 5% $H_2$).

An aliquot of 200 μL of each glycerol stock was inoculated into 20 mL of pre-reduced BG medium

(70% Bryant and Burkey Medium + 30% modified Gifu Anaerobic Medium, both purchased from Shandong Tuopu Biol-engineering Co., LTD.) and incubated for 24 hours at 37 °C in an anaerobic chamber. 1.5 mL of the culture was then allocated to each well of a 96-well plate. Each culture sample was divided into three groups including B (fecal bacteria with bile salt mixture), DB (fecal bacteria with addition of $10^7$ CFU of DSV/mL and bile salt mixture) and KB (heat-killed fecal bacteria with bile salt mixture). Besides KB as a negative control, an abiotic BG medium (M) with bile salt mixture added in was served as another negative control, which included three replicates. The number of DSV added was determined according to the reported relative abundance of *Desulfovibrio* species in healthy human beings, which is around 0.0001% - 1%[3]. 1% was chosen in this experiment to ensure a relative high number of DSV could be added. The specific CFU number was then determined based on the number of bacteria in a healthy human fecal sample. Briefly, 200 μL of one fecal glycerol stock was inoculated in 200 mL of BG medium and incubated at 37 °C for 24h in an anaerobic chamber. 1 mL of the culture was taken out and centrifuged at 10,000 rpm for 3 minutes. Medium was discarded and resuspended in 1 mL of 70% isopropanol for 30 minutes at room temperature to kill and fix the bacterial cells. Isopropanol was discarded after centrifugation at 10,000 rpm for 3 minutes and resuspended in 1 mL of 0.85% saline. The sample was diluted 100 times and was divided into halves, one served as the non-stained control and the other one was added with 0.75 μL of 20 mM propidium iodide (PI) to stain the bacterial DNA. Flow cytometry was then performed on the BD Accuri C6 Plus Flow Cytometer (Becton, Dickinson and Company, New Jersey, USA). The count of bacterial cells was around $10^9$ CFU/mL (Fig. S18). Therefore, the number of DSV added was $10^7$ CFU/mL.

Bile salt mixture contained 0.005mg/mL of sodium salt of taurocholic acid (TCA), 0.0018 mg/mL sodium salt of taurochenodeoxycholic acid (TCDCA), 0.0039 mg/mL sodium salt of glycocholic acid

(GCA) and 0.0032 mg/mL sodium salt of glycochenodeoxycholic acid (GCDCA), which was one tenth of the concentrations reported in the research of Huang's group[4] since the fecal sample culture only contained around $10^9$ CFU/mL of bacteria after 24 hours of incubation, around one tenth of the number of normal human gut microbes which is around $10^{10}$ - $10^{11}$ CFU[5]. After adding DSV or/and bile salts, the samples were allowed to incubate for another 24 hours. 1 mL of each sample was collected for 16s rRNA gene sequencing and the left 500 μL was collected for BA analysis.

***Quantification of BAs by UPLC-MS-MS***

50 mg of cecal content of germ-free mice from the 4th animal experiment or 100 μL of each bacteria culture from the *ex vivo* experiment was mixed with 400 μL acetonitrile and 0.1mm zirconium oxide beads and mixed thoroughly using a homogenizer (KZ-5F-3D, Wuhan Servicebio Technology Co., Ltd.). The samples were then centrifuged at 12,000 rpm at 4 °C for 10 minutes. 300 - 400 μL of the supernatant were pipetted into a 1.5 mL Eppendorf tube and dried using a vacuum centrifuge concentrator (Shanghai Jingxin Industrial Development Co., Ltd.). The sample pellet was resuspended in 100 - 200 μL methanol, vortexed for 1 minute, and centrifuged at 12,000 rpm at 4 °C for 10 minutes. 100 μL was pipetted from the top and run on a ExionLC AC HPLC-QTRAP5500 system (AB Sciex LLC, MA, USA).

BEH C18 column (1.7μm, 2.1 × 100mm) was used, column temperature was set at 35 °C and the injection volume was 2 μL. Mobile phase A was water with 0.01% formic acid and mobile phase B was acetonitrile with 0.01% formic acid. A gradient elution was set (20% B at 0-1 min, 35% B at 2.5 min, 70% B at 10 min, 99% B at 1 -12.5 min, 20% B at 12.6-15 min). Flow rate was 300 μL/min. The samples were placed at 4 °C during the whole process. Following mass spectrometry was performed in negative mode. The source conditions were set as follows: Source temperature: 550 °C; GS1: 60; GS2: 60; CUR: 35; IS: -4500 V.

MRM data was analyzed by Analyst software (v.1.7) to generate the quantitative data of each BA detected.

***Detection of amino acids in the chow diet using HPLC***

Amino acids were extracted by acid hydrolysis of the chow diet and detected by HPLC with automated OPA-derivatization (Agilent 1100, Agilent Technologies Inc., CA, USA). Tryptophan was not determined because acid hydrolysis destroys tryptophan. Briefly, 100 grams of the animal feed was weighed and 8 mL 6M HCl was added in. The sample was then concentrated using nitrogen blowing for 3 minutes and left for hydrolysis at 120 °C for 24 hours. Neutralization of the sample was done by adding 4.8 mL 10M NaOH and distilled water was added up to 25 mL. The sample was then filtered and centrifuged at 10,000 rpm for 10 minutes. 400 μL of the supernatant was added into a vial for the following HPLC analysis. The experiment was conducted in triplicate. 1 nmol/μL of 17 amino acid standards including lysine, methionine, cystine, arginine, histidine, phenylalanine, tyrosine, threonine, leucine, isoleucine, valine, aspartic acid, alanine, glutamic acid, glycine, proline and serine were prepared and standard curves were created. Agilent Hypersil ODS column (5 μm, 4.0mm × 250 mm) was used and column temperature was 40 °C. Flow rate was 1.0 mL/min. Mobile phase A was 27.6 mmol/L sodium acetate-triethylamine-tetrahydrofuran (v/v/v: 500 : 0.11: 2.5) and mobile phase B was 80.9 mmol/L sodium acetate-methanol-acetonitrile (v/v/v: 1 : 2 : 2). Gradient elution was set (8% B at 0 min, 50% B at 17 min, 100% B at 20.1 min, 0% B at 24 min). VWD was set at 338 nm for amino acid detection except proline which was detected at 262 nm.

References

1. Subramanian, A. et al. Gene set enrichment analysis: a knowledge-based approach for interpreting genome-wide expression profiles. *Proc Natl Acad Sci U S A* **102**, 15545-15550 (2005).
2. Javdan, B. et al. Personalized Mapping of Drug Metabolism by the Human Gut Microbiome. *Cell* **181**, 1661-1679 e1622 (2020).
3. Xiao, L. et al. A catalog of the mouse gut metagenome. *Nat Biotechnol* **33**, 1103-1108 (2015).
4. Shalon, D. et al. Profiling the human intestinal environment under physiological conditions. *Nature* **617**, 581-591 (2023).
5. Kastl, A.J., Jr., Terry, N.A., Wu, G.D. & Albenberg, L.G. The Structure and Function of the Human Small Intestinal Microbiota: Current Understanding and Future Directions. *Cell Mol Gastroenterol Hepatol* **9**, 33-45 (2020).

### Supplementary tables

Table S1 Nutrition facts of the animal diets

| | Chow diet | NRC95 purified diet | SAAL diet |
|---|---|---|---|
| Major ingredients | Fishmeal, wheat, corn, soybean meal, wheat bran, vitamin and mineral premix | Corn starch, corn maltodextrin, individual amino acids, soybean oil, cellulose, vitamins, and minerals | Corn starch, corn maltodextrin, individual amino acids, soybean oil, cellulose, vitamins, and minerals |
| Calories/kg | 3520 | 3800 | 3800 |
| Water content (%) | 9.2 | 6.6 | 6.6 |
| Fat (%) | 5.3 | 7 | 7 |
| Carbohydrates (%) | 57.2 | 64.3 | 65.3 |
| Protein (%) | 22.1 | 17.8 | 16.8 |
| Ash (%) | 6 | 4.17 | 4.17 |
| Lysine (g/kg) | 13.1 | 13 | 13 |
| Methionine (g/kg) | 5.2 | 8.2 | 1.5 |
| Cystine (g/kg) | 0.7 | 3.5 | 0 |
| Arginine (g/kg) | 11.5 | 6.4 | 6.4 |
| Histidine (g/kg) | 2.5 | 4.6 | 4.6 |
| Tryptophan (g/kg) | ≥ 2.5 | 2.1 | 2.1 |
| Phenylalanine (g/kg) | 9.2 | 8.8 | 8.8 |

| | | | |
|---|---|---|---|
| Tyrosine (g/kg) | 4.6 | 9.3 | 9.3 |
| Threonine (g/kg) | 6.5 | 6.7 | 6.7 |
| Leucine (g/kg) | 14.8 | 15.4 | 15.4 |
| Isoleucine (g/kg) | 8.3 | 8.5 | 8.5 |
| Valine (g/kg) | 10.1 | 10 | 10 |
| Aspartic acid (g/kg) | 19.8 | 12.2 | 12.2 |
| Alanine (g/kg) | 9.2 | 4.6 | 4.6 |
| Glutamic acid (g/kg) | 41.5 | 36.3 | 36.3 |
| Glycine (g/kg) | 9.0 | 3.2 | 3.2 |
| Proline (g/kg) | 13.3 | 20.5 | 20.5 |
| Serine (g/kg) | 8.4 | 9.7 | 9.7 |
| Calcium (ppm) | 10000 - 18000 | 5000 | 5000 |
| Phosphorus (ppm) | 6000 - 12000 | 3000 | 3000 |
| Potassium (ppm) | ≥ 5000 | 3600 | 3600 |
| Sodium (ppm) | ≥ 2000 | 1039 | 1039 |
| Magnesium (ppm) | ≥ 2000 | 513 | 513 |
| Iron (ppm) | ≥ 100 | 45 | 45 |
| Zinc (ppm) | ≥ 30 | 38 | 38 |
| Manganese (ppm) | ≥ 75 | 10 | 10 |
| Copper (ppm) | ≥ 10 | 6 | 6 |
| Iodine (ppm) | ≥ 0.5 | 0.2 | 0.2 |
| Chromium (ppm) | NA | 1 | 1 |

| | | | |
|---|---|---|---|
| Sulfur (ppm) | NA | 300 | 300 |
| Chloride (ppm) | NA | 1631 | 1631 |
| Vitamin A (IU/g) | ≥ 14 | 4 | 4 |
| Vitamin D (IU/g) | ≥ 1.5 | 1 | 1 |
| Vitamin E (IU/g) | ≥ 0.12 | 0.075 | 0.075 |
| Vitamin K (ppm) | ≥ 5 | 0.9 | 0.9 |
| Thiamine (ppm) | ≥ 13 | 5 | 5 |
| Riboflavin (ppm) | ≥ 12 | 6 | 6 |
| Niacin (ppm) | ≥ 60 | 30 | 30 |
| Pantothenic acid (ppm) | ≥ 24 | 15 | 15 |
| Vitamin B6 (ppm) | ≥ 12 | 6 | 6 |
| Choline (ppm) | ≥ 1250 | 1000 | 1000 |
| Folate (ppm) | ≥ 6 | 2 | 2 |
| Biotin (ppm) | ≥ 0.2 | 0.2 | 0.2 |
| Vitamin B12 (ppb) | ≥ 22 | 25 | 25 |

NA: not available.

Nutrient contents in the chow diet except amino acids are referred to Chinese standard GB 14924.3-2010 for Laboratory animals – Nutrients for formula feed, which only provides a range or minimal content of each nutrient.

Amino acids were extracted by acid hydrolysis of the chow diet and detected by HPLC with automated OPA-derivatization (Agilent 1100, Agilent Technologies Inc., CA, USA). Specific methods can be found in Supplementary methods. Tryptophan was not determined because acid hydrolysis destroys tryptophan. Recommended content of tryptophan is referred to Chinese standard GB 14924.3-2010 for Laboratory animals – Nutrients for formula feed.

Table S2 Primer list

| Gene name | Primers |
| --- | --- |
| *Cyp7a1* | F: AGCAACTAAACAACCTGCCAGTACTA |
| | R: GTCCGGATATTCAAGGATGCA |
| *Cyp8b1* | F: CATGAAGGCTGTGCGTGAGGAA |
| | R: CATCACGCTGTCCAACACTGGA |
| *Nr1h4* | F: TCCAGGGTTTCAGACACTGG |
| | R: GCCGAACGAAGAAACATGG |
| *Sod1* | F: AGCATGGCGATGAAAGCGG |
| | R: CCTGCACTGGTACAGCCTTGT |
| *Sod2* | F: AACGCCACCGAGGAGAAGTA |
| | R: TCCAGCAACTCTCCTTTGGGT |
| *Gsta2* | F: ACCGTTACTTGCCTGCCTTT |
| | R: GCTGGCATCAAGCTCTTCAAC |
| *Gstm1* | F: AAAGCACCACCTGGATGGAGA |
| | R: GCTCCCCAGCAAAGGGTTT |
| *Gstm2* | F: TGGACTTTCCCAATCTGCCCT |
| | R: TCATCTTCTCAGGGAGACCCTC |
| *Gstp1* | F: TCTACGCAGCACTGAATCCG |
| | R: GGAGCTGCCCATACAGACAA |
| *Zo1* | F: AGGACACCAAAGCATGTGAG |
| | R: GGCATTCCTGCTGGTTACA |

| | |
|---|---|
| *Ocln* | F: CGGTACAGCAGCAATGGTAA |
| | R: CTCCCCACCTGTCGTGTAGT |
| *Aoah* | F: GTTTTCCCAACGCTGCGGGG |
| | R: TGGCCTTCTGCCCGGGTACA |
| *Gapdh* | F: AGGTCGGTGTGAACGGATTTG |
| | R: TGTAGACCATGTAGTTGAGGTCA |

### Supplementary figures

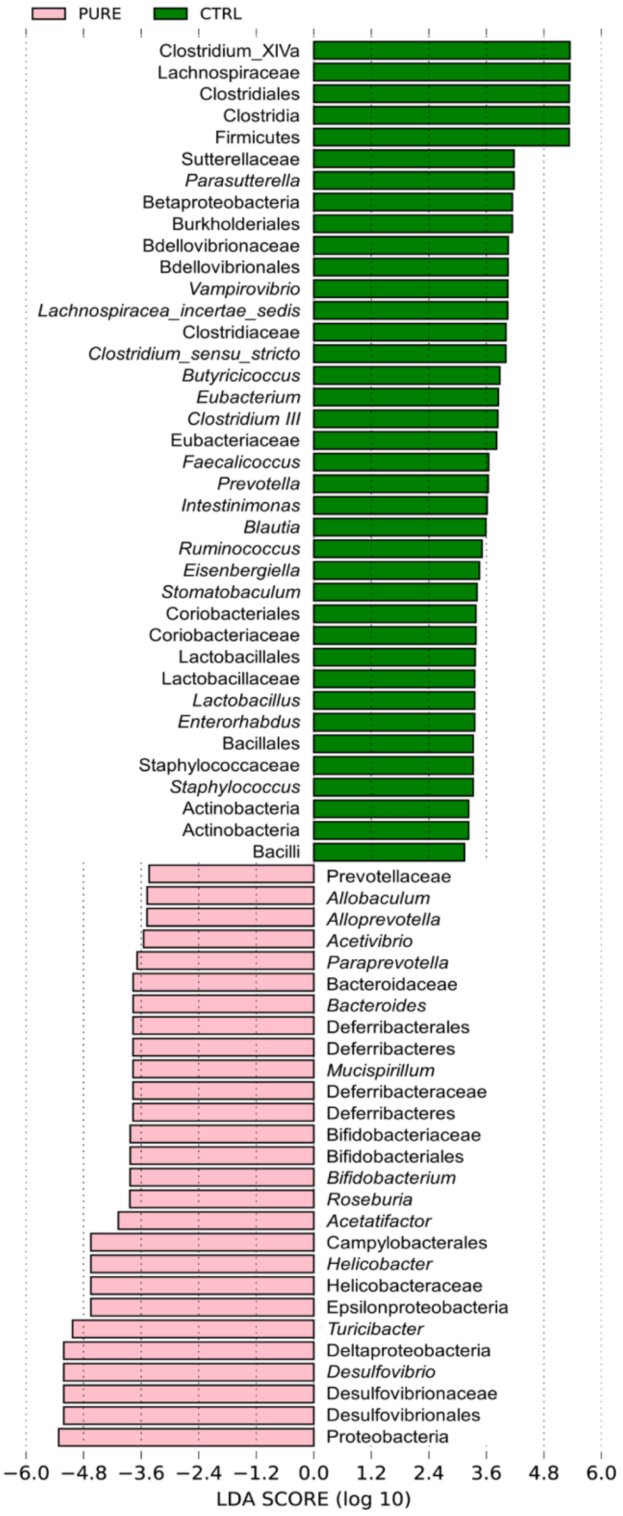


Fig. S1. LDA score of LEfSe for differential microbial biomarkers in mice fed with the chow and purified diet. OTUs with LDA score >2 were screened and represented the microbial biomarkers in that group. n=10 mice per group.

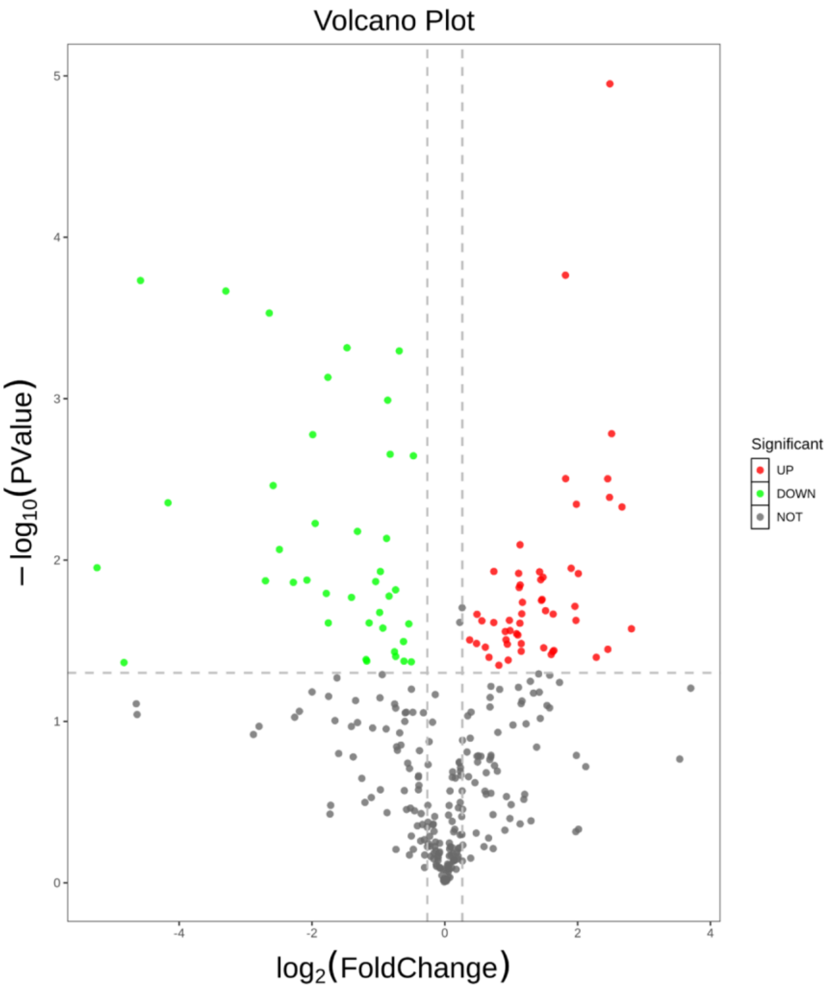


Fig. S2. Volcano plot of 348 metabolites detected in the cecal content of the CTRL and PURE group. Red dots represent metabolites up-regulated in the PURE group and green dots represent metabolites down-regulated in the PURE group. n=3 mice per group.

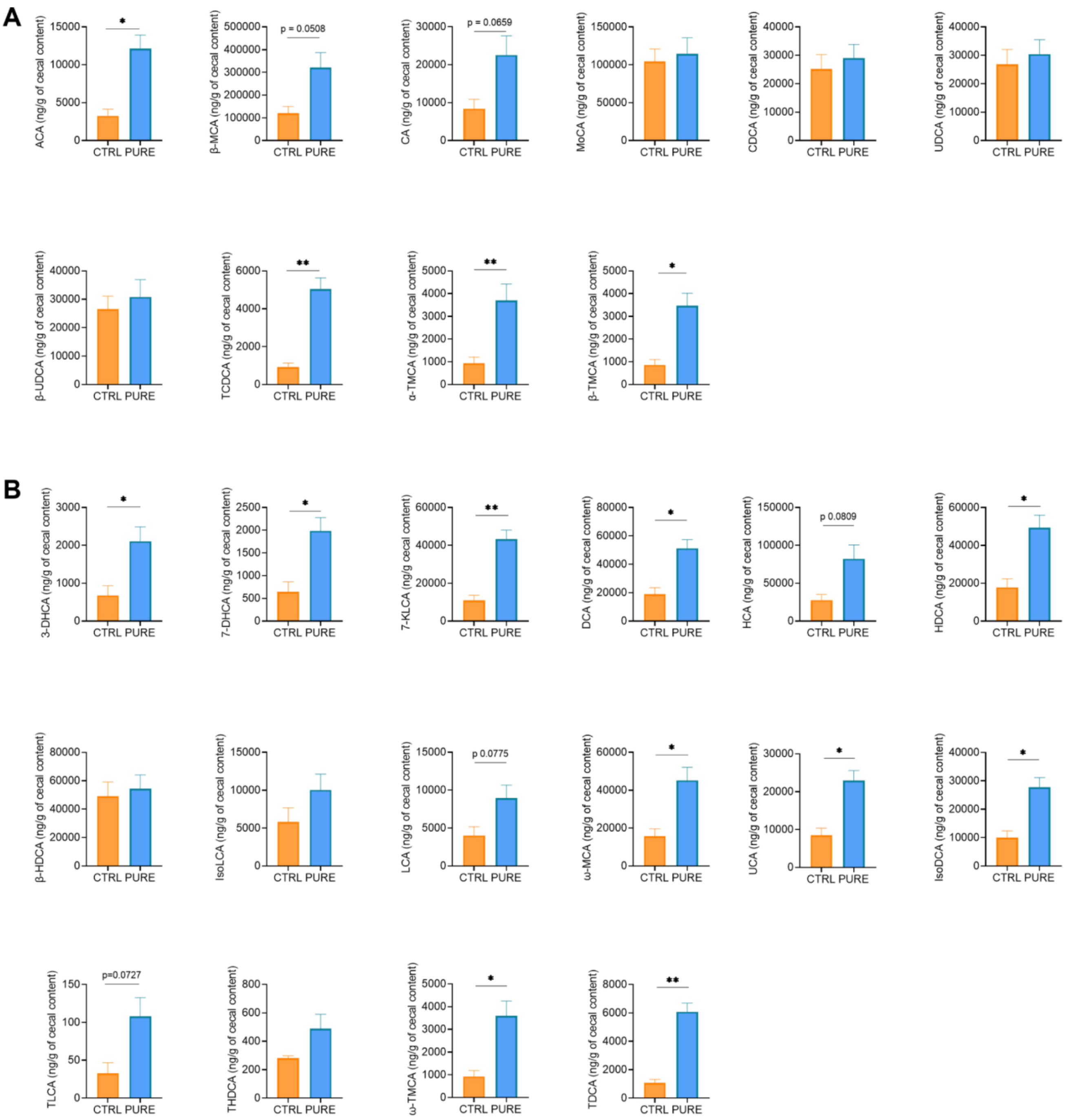


Fig. S3. Concentrations of 26 types of BAs in the CTRL and PURE group. The BAs with data points less than 3 in any of the two groups were not included. (A) Concentrations of all detected primary BAs. (B) Concentrations of all detected secondary BAs. n=3 mice per group. Data are represented as the mean ± SEM, two-tailed Student t-test or Mann-Whitney test, *$p < 0.05$, **$p < 0.01$. ACA: apocholic acid, β-MCA: beta-muricholic acid, CA: cholic acid, TCDCA: taurochenodeoxycholic acid, β-UDCA: beta-ursodeoxycholic acid, UDCA: ursodeoxycholic acid, CDCA: chenodeoxycholic acid, MoCA: murocholic acid, α-TMCA: tauro-α-muricholic acid, β-TMCA: tauro-β-muricholic acid, 7-DHCA: 7-dehydrocholic acid, 7-KLCA: 7-ketolithocholic acid, DCA: deoxycholic acid, 3-DHCA: 3-dehydrocholic acid, HCA: hyocholic acid, HDCA: hyodeoxycholic acid, β-HDCA: beta-hyodeoxycholic acid, IsoDCA: isodeoxycholic acid, IsoLCA: isolithocholic acid, LCA: lithocholic acid, ω-MCA: omega-muricholic acid, GHDCA: glycohyodeoxycholic acid, TDCA: taurodeoxycholic acid, TLCA: taurolithocholic acid, THDCA: taurohyodeoxycholic acid, ω-TMCA: tauro-ω-muricholic acid.

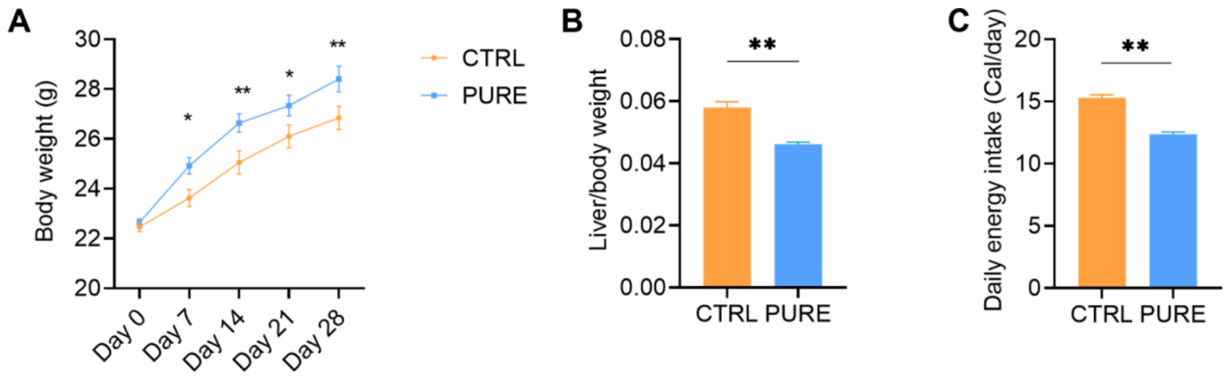


Fig. S4. Physiological changes of mice fed with chow and purified diet. (A) Body weight change during 28 days. (B) Liver/body weight ratio. (C) Daily energy intake. n = 10 mice per group. All data are represented as mean ± SEM, two-tailed Student t-test, *$p < 0.05$, **$p < 0.01$.

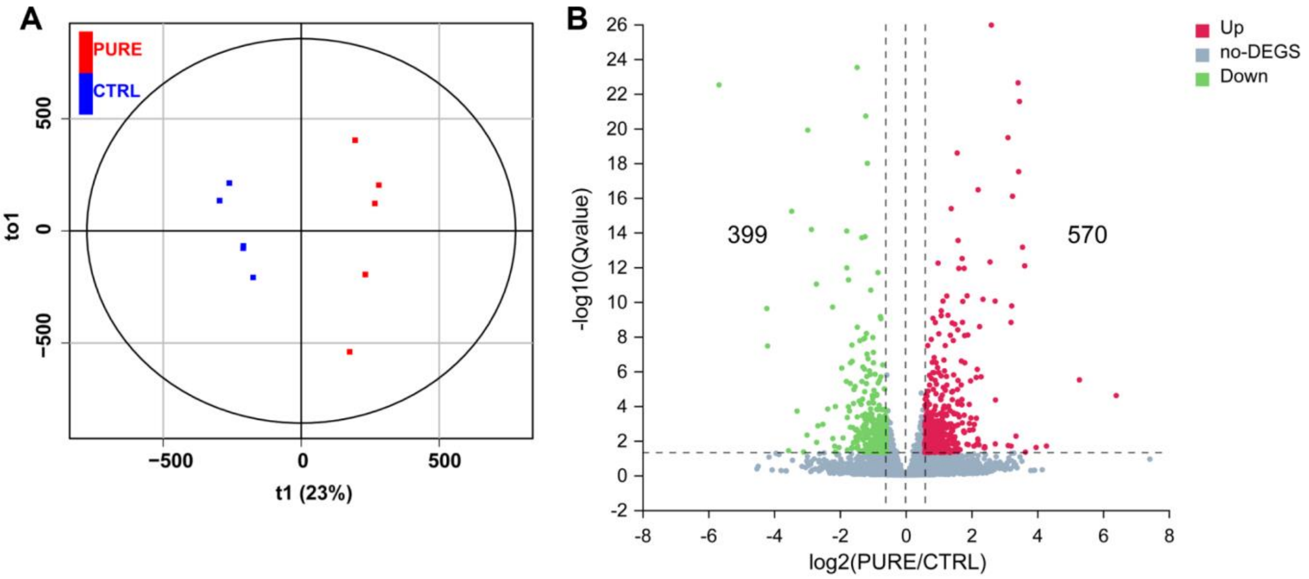


Fig. S5. Different liver gene expression patterns in mice fed with chow and purified diet using transcriptomic analysis. (A) OPLS-DA score. (B) Volcano plot of gene expression levels. Gene expressions with log2(PURE/CTRL) > 0.58 and Qvalue (adjusted p value) < 0.05 are marked red and gene expressions with log2 (PURE/CTRL) < 0.58 and Qvalue (adjusted p value) < 0.05 are marked green. n = 5 mice per group.

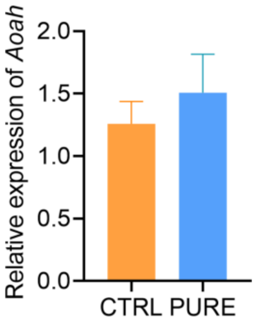


Fig. S6. Relative expression of hepatic *Aoah* in the CTRL and PURE group. There was no significant difference between these two groups. n = 4-5 mice per group. All data are represented as mean ± SEM, two-tailed Student t-test.

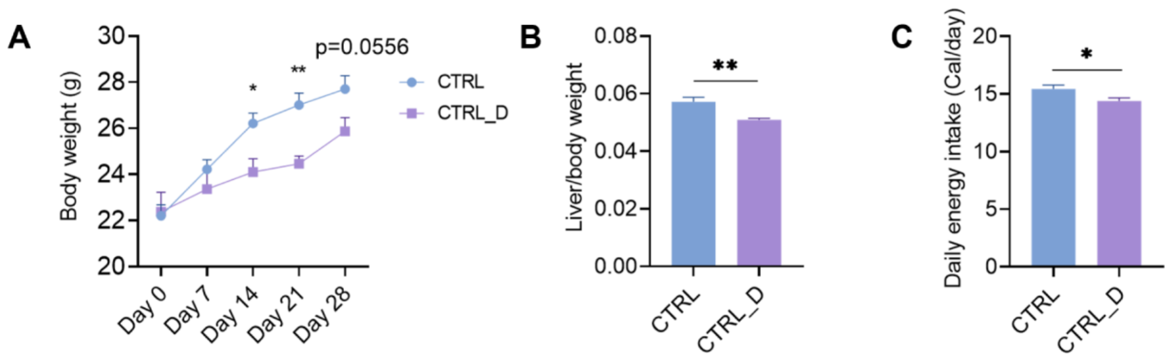


Fig. S7. Physiological changes of mice in the CTRL and CTRL_D group. (A) Body weight change during 28 days. (B) Liver/body weight ratio. (C) Daily energy intake. n = 5 mice per group. All data are represented as mean ± SEM, two-tailed Student t-test, *$p < 0.05$, **$p < 0.01$.

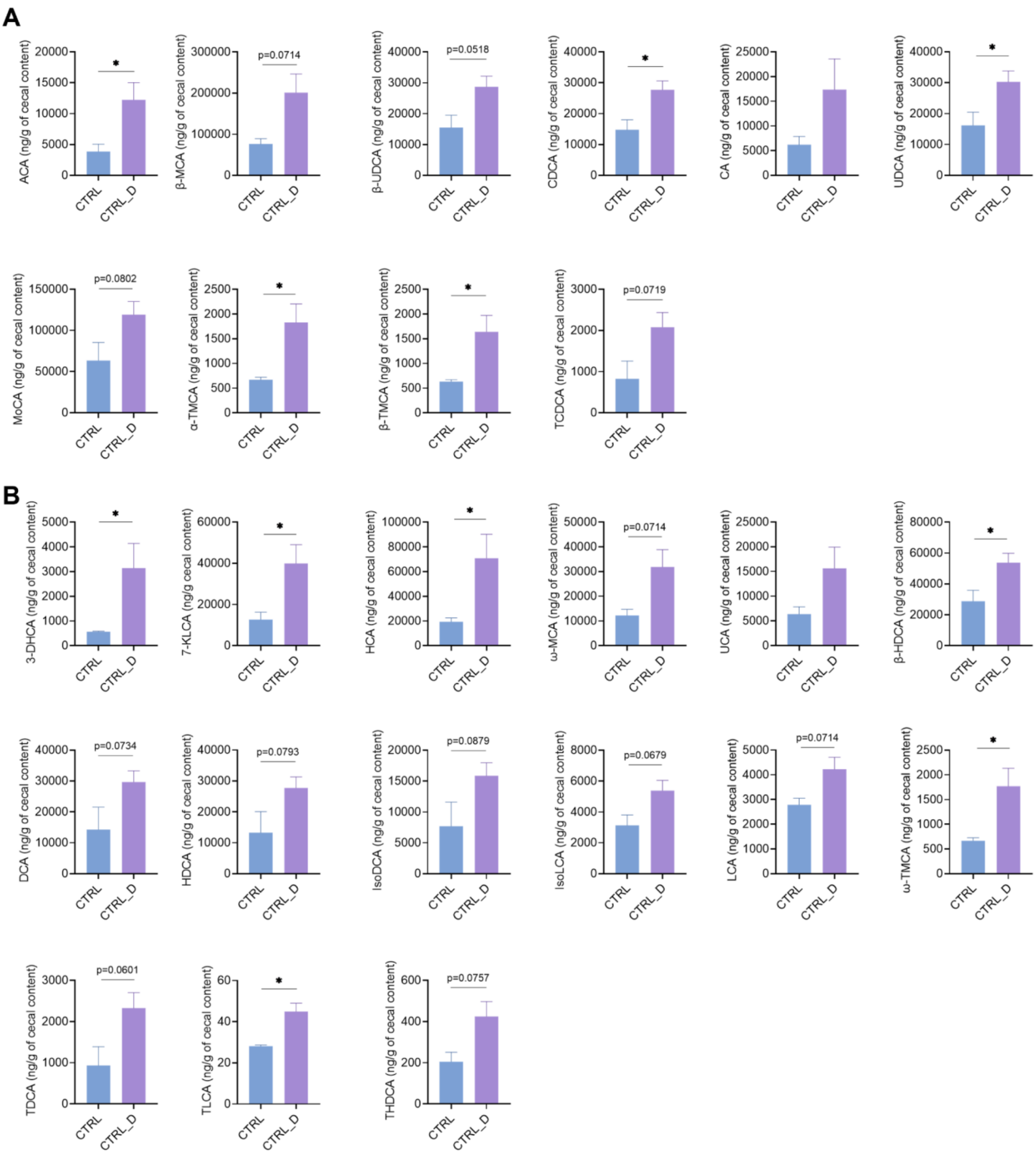


Fig. S8. Concentrations of 24 types of BAs in the CTRL and CTRL_D group. The BAs with data points less than 3 in any of the two groups were not shown. (A) Concentrations of all detected primary BAs. (B) Concentrations of all detected secondary BAs. n = 3-5 mice per group. All data are represented as the mean ± SEM, two-tailed Student t-test or Mann-Whitney test, *$p < 0.05$.

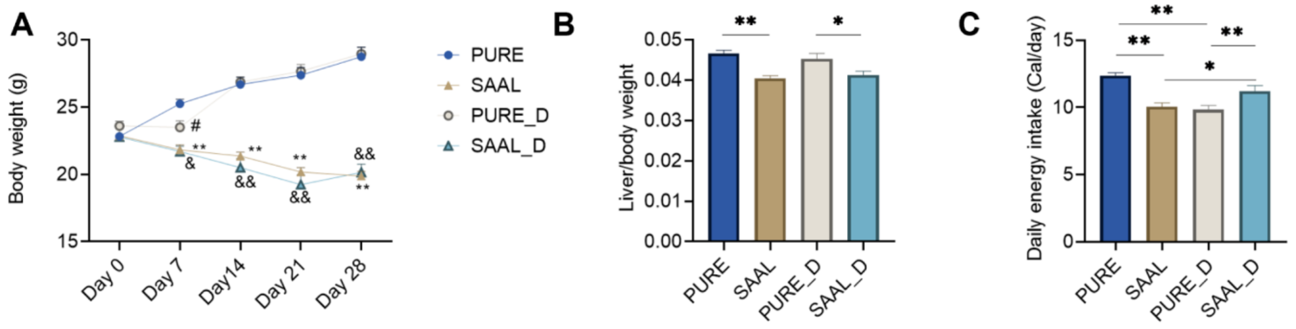


Fig. S9. Physiological changes of mice in the CTRL and CTRL_D group. (A) Body weight change during 28 days. (B) Liver/body weight ratio. (C) Daily energy intake. n = 6 mice per group. All data are represented as mean ± SEM, one-way ANOVA with Holm-Sidak post hoc test, *$p < 0.05$, **$p < 0.01$.

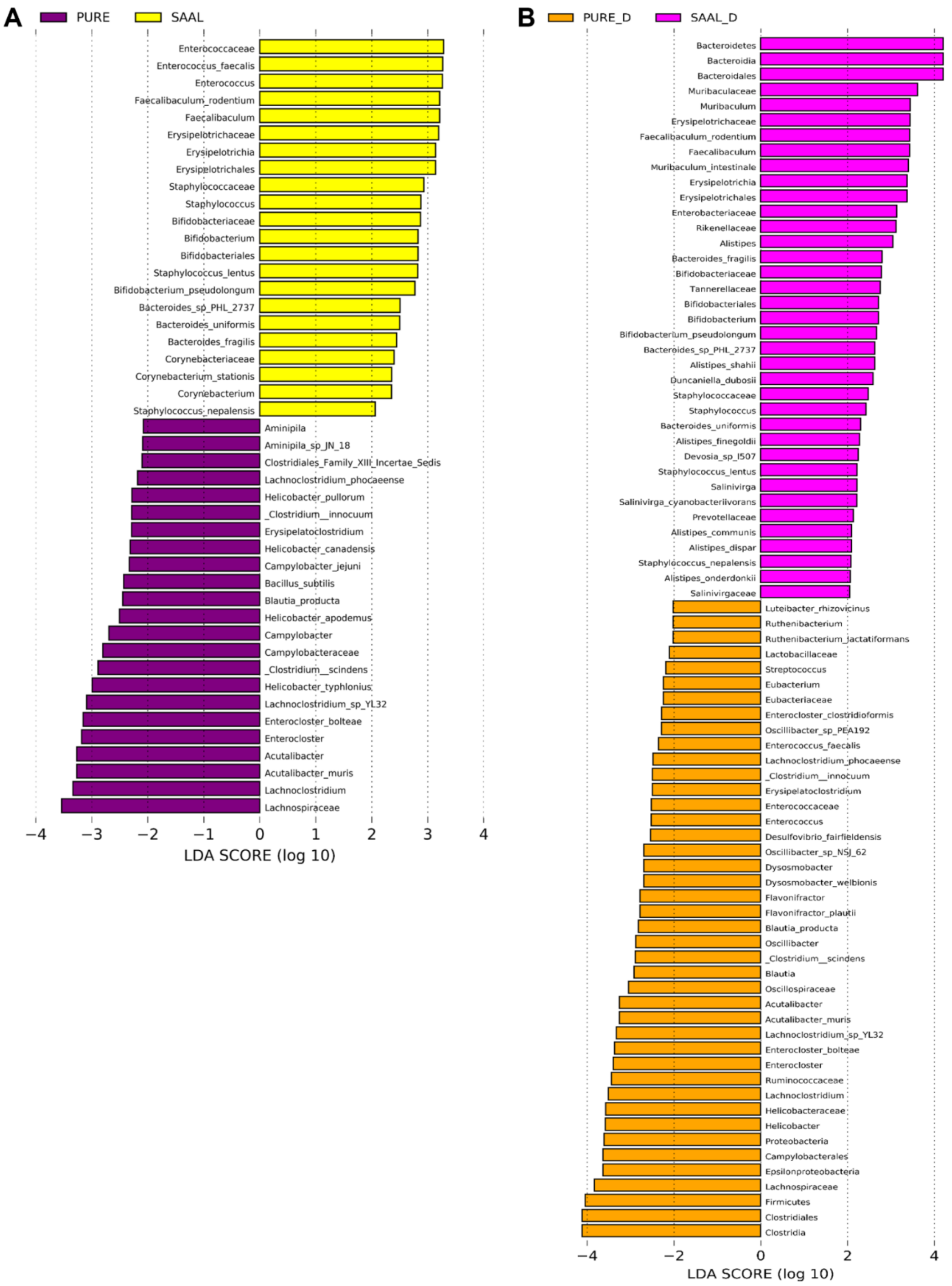


Fig. S10. LDA score for differential microbial biomarkers in LEfSe between mice fed with the purified diet and SAAL diet. (A) LDA score between the PURE and SAAL group. (B) LDA score between the

PURE_D and SAAL_D group. OTUs with LDA score >2 were screened and represented the microbial biomarkers in that group. n = 5 mice per group.

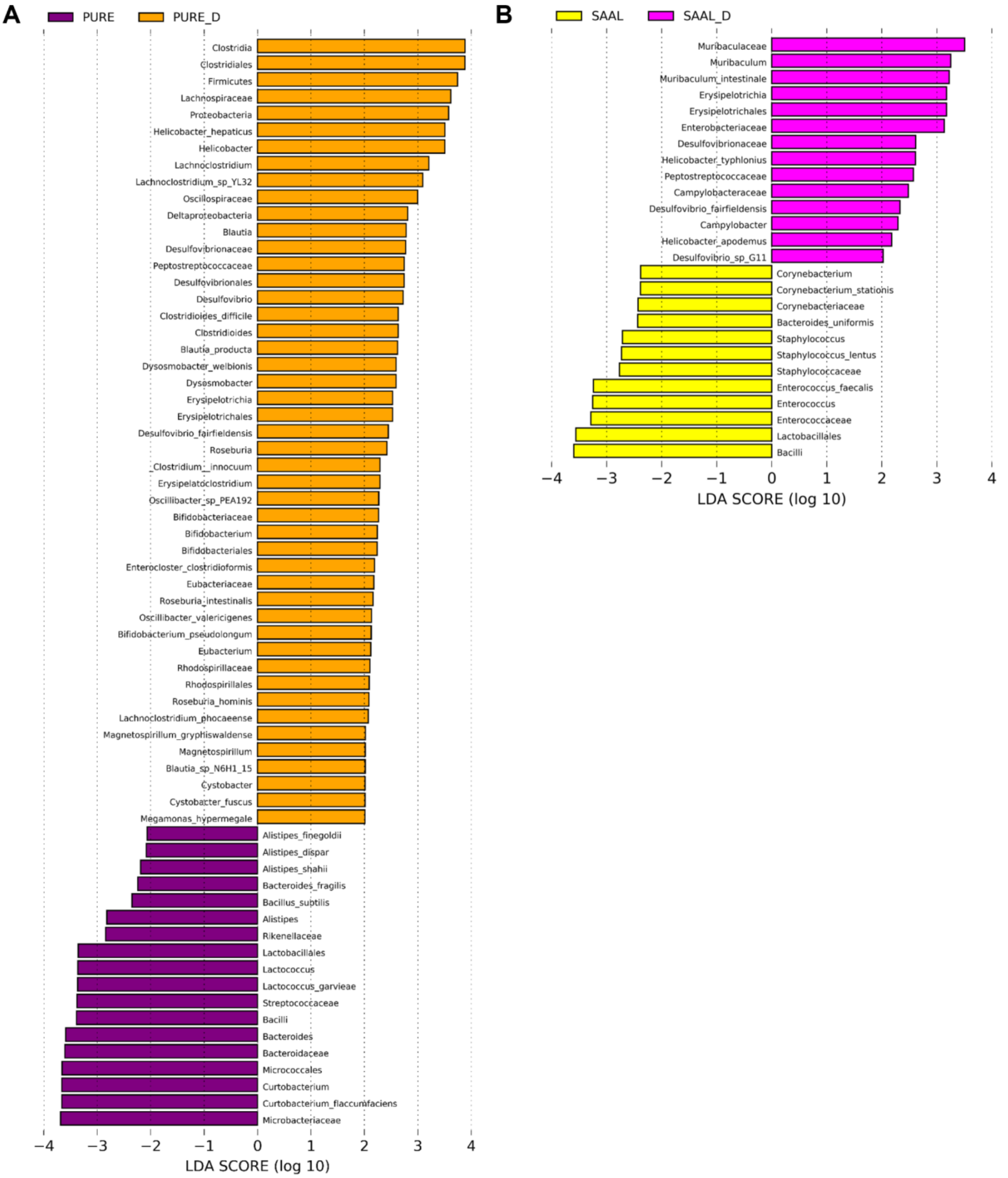


Fig. S11. LDA score for differential microbial biomarkers in LEfSe between mice with or without DSV administration. (A) LDA score between the PURE_D and PURE group. (B) LDA score between the SAAL_D and SAAL group. OTUs with LDA score >2 were screened and represented the microbial biomarkers in that group. n =5 mice per group.

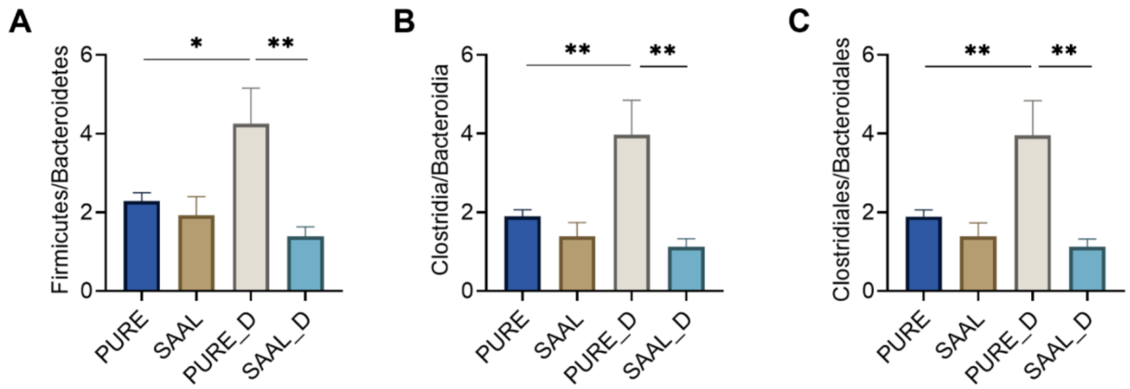


Fig. S12. Firmicutes/Bacteroidetes ratio, Clostridia/ Bacteroidia ratio, and Clostridiales/Bacteroidales ratio in the mouse gut of PURE, SAAL, PURE_D and SAAL_D group. (A) Firmicutes/Bacteroidetes. (B) Clostridia/Bacterodia. (C) Clostridiales/Bacteroidales. n = 5 mice per group. All data are represented as the mean ± SEM, one-way ANOVA with FDR post hoc test (Two-stage step-up method of Benjamini, Krieger and Yekutieli), $*p < 0.05$, $**p < 0.01$.

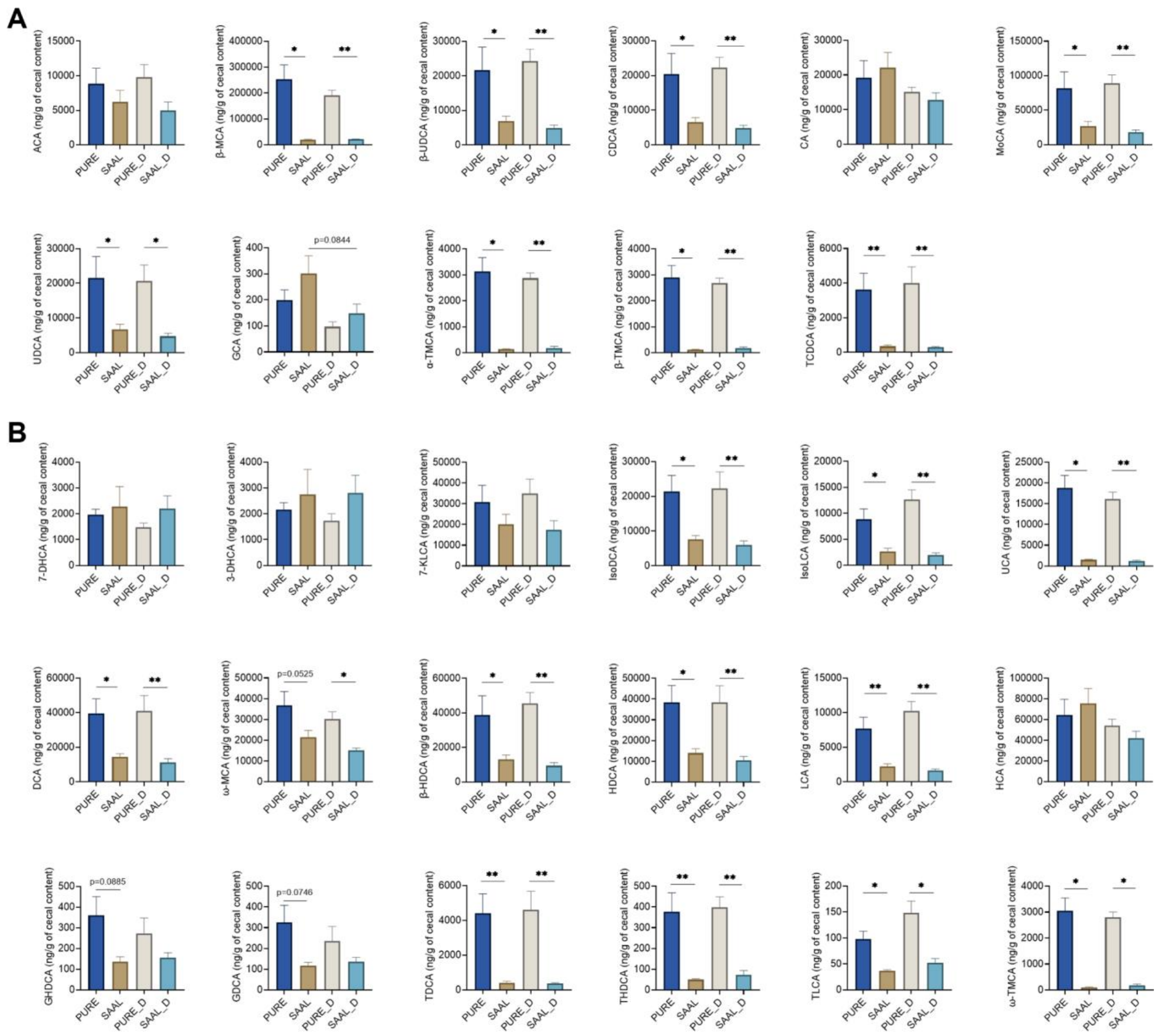

Fig. S13. Concentrations of 29 types of BAs in the PURE, SAAL, PURE_D and SAAL_D group. The BAs with data points less than 3 in any of the groups were not shown. (A) Concentrations of all detected primary BAs. (B) Concentrations of all detected secondary BAs. n = 5-6 mice per group. All data are represented as the mean ± SEM, one-way ANOVA or Kruskal-Wallis test with FDR post hoc test (Two-stage step-up method of Benjamini, Krieger and Yekutieli), $*p < 0.05$, $**p < 0.01$.

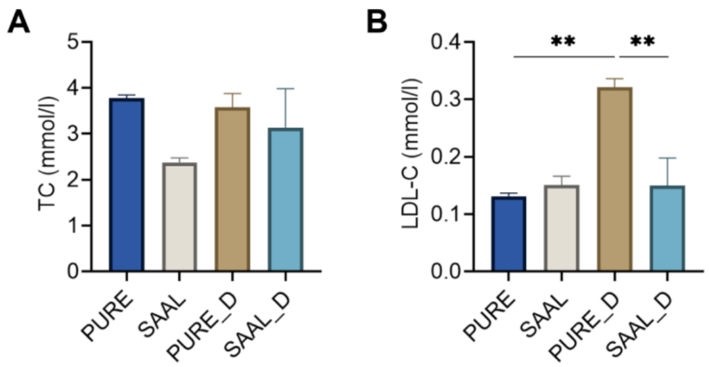


Fig. S14. Serum cholesterol level of mice in the PURE, SAAL, PURE_D and SAAL_D group. (A) Concentration of total cholesterol (TC). (B) Concentration of low-density lipoprotein cholesterol (LDL-C). n = 3 mice per group. All data are represented as the mean ± SEM, one-way ANOVA with FDR post hoc test (Two-stage step-up method of Benjamini, Krieger and Yekutieli), $^{**}p < 0.01$.

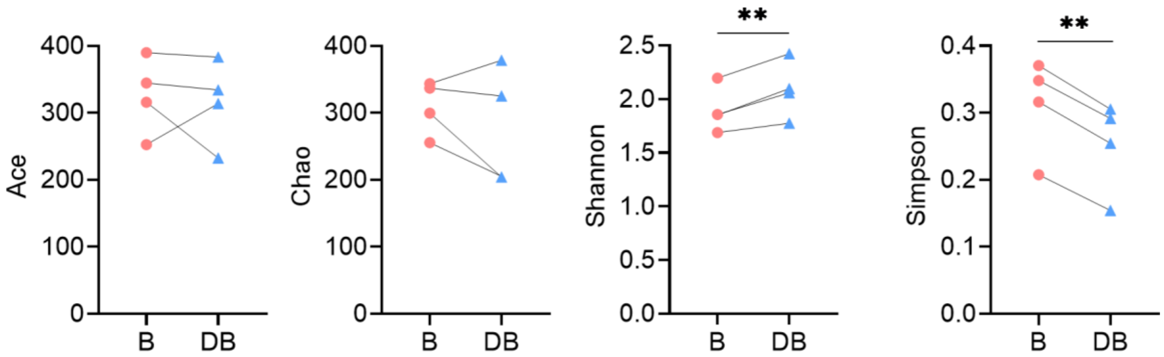


Fig. S15. Alpha diversity of fecal bacteria including Chao, Ace, Shannon and Simpson index from 4 healthy volunteers fermented with four primary BAs after 24h. n = 4. All data are represented as the mean ± SEM, one-tailed paired t-test or Wilcoxon matched-pairs signed rank test, $^{**}p < 0.01$.

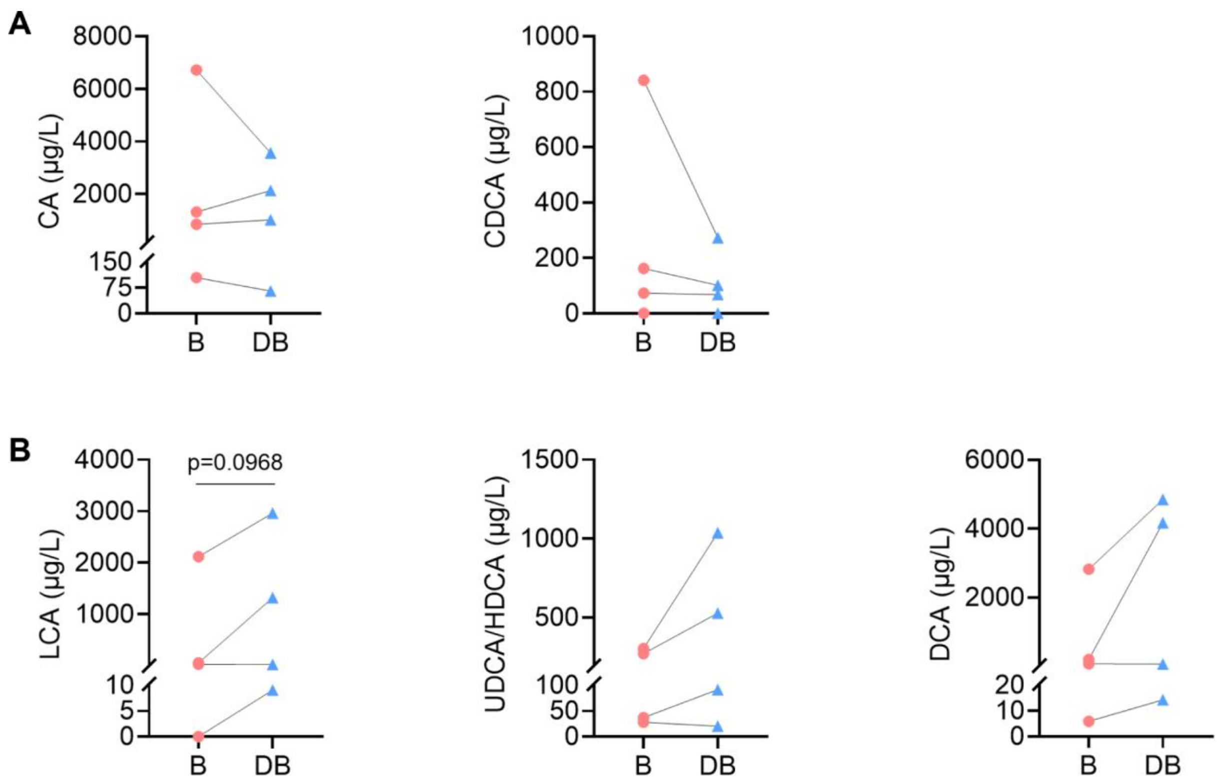


Fig. S16. Concentration of primary and secondary BAs detected in the fecal bacterial culture of 4 healthy volunteers. (A) Primary BAs including CA and CDCA. (B) Secondary BAs including LCA, UDCA/HDCA and DCA. UDCA and HDCA were put together because they cannot be separated due to very close elution time. n = 4. All data are represented as the mean ± SEM, one-tailed paired t-test or Wilcoxon matched-pairs signed rank test.

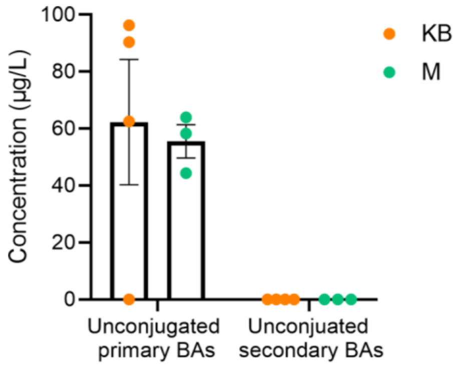


Fig. S17. Concentration of primary and secondary BAs in the heat-killed fecal bacterial culture and abiotic medium. n = 3-4. All data are represented as the mean ± SEM, two-tailed Student t-test.

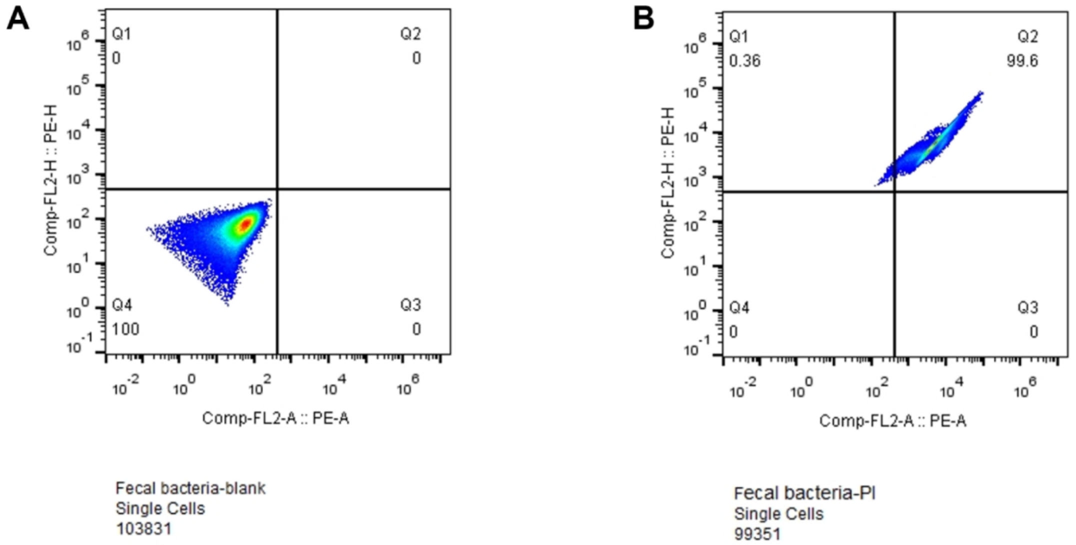


Fig. S18. Flow cytometry of fecal bacterial cells from one healthy volunteer after 24 h of incubation. (A) Negative control. The sample was diluted 100 times without propidium iodide (PI) staining. (B) PI-stained sample with 100 times of dilution. Volume for detection: 10 μL.
